# Knockout of Sorbs2 in Cardiomyocytes Leads to Dilated Cardiomyopathy in Mice

**DOI:** 10.1101/2022.02.13.480093

**Authors:** Jared M. McLendon, Xiaoming Zhang, Daniel S. Matasic, Mohit Kumar, Olha M. Koval, Isabella M. Grumbach, Sakthivel Sadayappan, Barry London, Ryan L. Boudreau

## Abstract

2.

**Rationale:** Sorbs2 is a cardiomyocyte-enriched, cytoskeletal adaptor protein, and given some evidence for its dysregulated expression in failing hearts, there is growing interest in understanding its roles in cardiac biology and disease. While Sorbs2 global knockout mice display lethal cardiomyopathy with severe arrhythmias, the underlying mechanisms remain unclear, and whether this results from intrinsic loss of Sorbs2 in cardiomyocytes is unknown, as Sorbs2 is also well-expressed in the nervous system and vasculature. In addition, the potential relevance of Sorbs2 in human cardiomyopathy remains underexplored.

**Objective:** To characterize the effects and potential underlying mechanisms of cardiomyocyte-specific deletion of Sorbs2 on cardiac structure and function in mice, and to further examine Sorbs2 dysregulation in failing hearts and explore potential links between Sorbs2 genetic variations and human cardiovascular disease phenotypes.

**Methods and Results:** We report that myocardial Sorbs2 expression is consistently upregulated in humans with ischemic and idiopathic cardiomyopathies, and in experimental animal models of heart failure (HF). We generated mice with cardiomyocyte-specific loss of Sorbs2 (Sorbs2-cKO) and found early atrial and ventricular conduction abnormalities, despite unaltered expression of primary action potential ion channels and gap junction proteins. At mid-life, Sorbs2-cKO mice exhibit impaired cardiac contractility with cardiomyofibers failing to maintain adequate mechanical tension. As a result, these mice develop progressive diastolic and systolic dysfunction, enlarged cardiac chambers, and die with congestive HF at approximately one year of age. Comprehensive survey of potential underlying mechanisms revealed that Sorbs2-cKO hearts exhibit defective microtubule polymerization and compensatory upregulation of structural proteins desmin, vinculin, and tubulins. Finally, consistent with our observations in mice, we identified suggestive links between Sorbs2 genetic variants and related human cardiac phenotypes, including conduction abnormalities, atrial enlargement, and dilated cardiomyopathy.

**Conclusions:** Our studies show that Sorbs2 is essential for maintaining cytoskeletal structural integrity in cardiomyocytes likely through strengthening the interactions between microtubules and other structural proteins at crosslink sites. Overall, this study provides key insights into the critical role for Sorbs2 in cardiomyocytes and likely other cell types in maintaining normal cardiac structure and function and highlights its potential clinical relevance.

## 3. Introduction

Heart failure (HF) remains a leading cause of worldwide morbidity and mortality, due in part to multiple etiological origins of cardiomyopathy and ineffective treatments. Along with several well-characterized pathogenic molecular signaling cascades, cardiomyocyte structural and cytoskeletal adapter proteins are known to play a role in HF development and progression. Individual cardiomyocytes string together into myofibers through complex connections that organize intra- and inter-cellular cytoskeletons into continuous longitudinal structures via intercalated discs (ICDs) and costameres. ICDs are composed of diverse cell-cell connections including components of adherens junctions and desmosomes that link actin and desmin (Des) networks of adjacent cardiomyocytes into cohesive units, as well as gap junctions, which facilitate rapid intercellular electrical conduction and communication through aqueous pores.

Costameres attach cardiomyocytes to the extracellular matrix through lateral connections of the sarcomeres to the sarcolemma at z-discs through membrane proteins, cytoplasmic actin, desmin, various actin-associated proteins, integrins, and dystrophin complexes. Disruptions in these structures are found in diseased hearts showing abnormal conduction and contractile dysfunction. In addition, genome-wide linkage studies for cardiac conduction anomalies and dilated cardiomyopathy (DCM) point to several cytoskeletal structural genes common to the sarcomeres, z-discs, costameres, and ICDs, among others.^1^

Sorbs2 belongs to the SORBS family of adaptor proteins that facilitate protein-protein interactions among many cytoskeletal and membrane-associated proteins including actin, actinin, vinculin, and various signaling kinases. Sorbs2 is relatively broadly expressed across tissues, with highest and enriched expression in cardiac myocytes and smooth muscle containing tissues.^2, 3^ Prior reports indicate that Sorbs2 expression is altered in the setting of myocardial infarction and diabetic cardiomyopathy, and its suppression *in vitro* induces cardiomyocyte hypertrophy.^4–6^ In addition, recent *in vivo* studies suggest both pathogenic and protective roles for Sorbs2 in the development of arrhythmogenic right ventricular cardiomyopathy (ARVC)^7^ and left ventricular non-compaction (LVNC).^8^ Specifically, Ding et al. reported that global Sorbs2 knockout mice (KO) develop lethal ARVC with severe right ventricle (RV) failure at 4-6 months of age. Whereas, Li et al. found that Sorbs2 expression increased in LVNC and that AAV-mediated overexpression of Sorbs2 in mouse hearts caused cardiac hypertrophy and severe HF within 3 weeks.^8^ Despite clear interest in understanding the relevance of Sorbs2 in cardiac biology and disease, the cell-specific *in vivo* role for Sorbs2 in cardiomyocytes has not been examined using conditional KO mice. Also, whether myocardial Sorbs2 expression is broadly dysregulated across different cardiomyopathic states in patients and mouse models remains to be systematically evaluated. Herein, we address these knowledge gaps and report that Sorbs2 expression is consistently upregulated across a variety of cardiomyopathies in humans and rodents, and that cardiomyocyte-specific loss of Sorbs2 in mice is sufficient to cause adverse remodeling of myocardial cytoskeletal proteins. This leads to a progressive DCM phenotype encompassing cardiac conduction defects, systolic and diastolic dysfunction, depressed myofiber contractility, and ultimately congestive HF and premature death, representing an intriguingly distinct phenotype compared to global Sorbs2 KO mice.^7^ While assessing the potential translational relevance of these findings through query of available human GWAS data, we identified several common genetic variants in SORBS family genes that are significantly linked to decreased Sorbs2 expression, altered cardiac conduction and DCM phenotypes that are consistent with our observations in mice.

## 4. Methods

### 4.1 Generation of Cardiomyocyte Sorbs2 Knockout Mice

The University of Iowa Animal Care and Use Committee (IACUC) approved all animal experiments described in this manuscript. All rodents used in these studies were housed under 12/12-hour light/dark cycle with access to food and water ad libitum. The Sorbs2 floxed mice (Sorbs2 fl/fl) harboring loxP sites flanking Sorbs2 exon 12, were generated by Dr. G. Feng^9^ and acquired from (Jax Labs stock # 028600). These mice were intercrossed to C57BL/6J transgenic mice containing the α-myosin heavy chain promoter-driven Cre recombinase (αMHC-Cre),^10^ provided by Dr. C. Grueter,^11^ to generate Sorbs2 cardiac-specific knockout (cKO) mice defined as αMHC-Cre positive Sorbs2 fl/fl mice. Most studies used Cre-negative Sorbs2 fl/fl littermates as controls, and some also included transgenic αMHC-Cre only mice as additional controls. Both male and female mice were used. Raw data from physiology experiments were analyzed by investigators blinded to animal genotypes.

### 4.2 Mouse Models of Cardiac Dysfunction

#### Viruses and delivery

A cDNA encoding constitutively-active CaMKII (T287D; gift from Dr. Mark Anderson) was cloned into a standard AAV2-ITR plasmid with a Cardiac Troponin-T (cTnT) promoter. AAV2/9 viruses were made by the University of Iowa Viral Vector Core. AAV2/9-CaMKII or AAV2/9-GFP control viruses were injected into 3-week-old C57/BL6J male mice anesthetized with isoflurane (2-3%, to effect) via intrajugular vein injection (dose for CaMKII = 3.5E+10 vg/g or GFP = 1.0E+10 vg/g), which results in high (>90%) and stable transduction of mouse cardiomyocytes. Mice were euthanized 3 weeks post injection.

#### Transverse Aortic Constriction (TAC)

Minimally invasive TAC was performed as previously described with few changes.^12^ At the level of the suprasternal notch, a partial sternotomy and thyroid retraction was used to visualize the aortic arch on 8-week-old C57BL/6J male mice anesthetized with ketamine/xylazine (100/10 mg/kg, IP). The aorta was isolated and then constricted with a titanium ligating clip (Teleflex, #005200) gapped on 38-gauge acupuncture needles, and placed between the right innominate and left common carotid arteries. Sham mice underwent the same procedure, but without constriction of the aorta. Mice were euthanized 8 weeks post TAC.

### 4.3 Echocardiography

Echocardiography was performed on conscious mice, with mild sedation (Midazolam 5 mg/kg, SC) restrained in the operator’s hand, using a Vevo2100 imaging system (VisualSonics, Toronto, Canada). Two-dimensional cine loops were acquired in both long- and short-axis planes to measure standard parameters of cardiac structure and function according to the endocardial and epicardial area protocol.

### 4.4 Electrocardiography (EKG)

Simultaneous three-lead EKG were acquired from anesthetized mice (2% isoflurane) on a heated platform using needle electrodes inserted subcutaneously in limbs (Indus, Rodent Surgical Monitor+ and analog output device). Data were analyzed using standard mouse settings, including all beats, with time selections of 10 seconds with 4 beat averaging for EKG analyses, and 60 seconds for HRV analyses (AD Instruments, Powerlab 8/8, Labchart Pro v8.1.13, EKG module v2.4, and HRV module v2.0.3).

### 4.5 LV Hemodynamics with Dobutamine (DOB) Challenge

Mice were anesthetized with 2% isoflurane and placed on a surgical monitoring board to collect EKG and maintain body heat (Indus, Rodent Surgical Monitor+). Hemodynamics were acquired from a pressure catheter and controller (Millar, SPR-671 and PCU-2000) inserted in the right common carotid artery and pushed to the left ventricle (LV). For stress tests, a fluid-filled catheter inserted in the left jugular vein was connected to a syringe pump to perform a step-by-step ramped Dobutamine (DOB) infusion maintained for 2 min at each step (dose range 2-12 ng/g/min).^13^ Data were analyzed using standard mouse settings without excluding “outlier” beats and binned into 10 second block averages from baseline throughout the dose response curve (AD Instruments, Powerlab 8/8, Labchart Pro v8.1.13, Blood Pressure module v1.4).

### 4.6 Myofiber Mechanics

Frozen heart tissues were thawed in a pCa 9.0 relaxing buffer **(Supp. Table 1)**, trimmed into fiber bundles (∼1 mm long) and skinned overnight in 1% (w/v) Triton X-100. Aluminum T-clips were attached to ends of straight fiber bundles, attached to a force transducer and length controller (Aurora Scientific Inc., Aurora, ON, Canada) and sarcomere length set to 2.1. The fibers were exposed to calcium (dose range pCa 9.0 to pCa 4.5) obtained by proportional mixing of relaxing and activating buffers **(Supp. Table 1),** as described previously.^14^ Fibers were discarded if they exhibited greater than 20% rundown in force over the experiment. Fiber integrity was tested by measuring maximal tension after the experiment (100% activation). Any fibers that did not maintain 80% or greater maximal tension were excluded from analysis.

Muscle dimensions (cross-sectional area, length) were determined using an ocular micrometer and used to normalize contractile force assuming elliptical fiber shape.

### 4.7 Western Blot

Frozen heart tissues were homogenized with a bead mill (Qiagen, TissueLyserII) in tissue lysis buffer **(Supp. Table 1)**. Homogenates were sonicated, clarified by centrifugation (16,000×g, 10 mins, 4℃), and normalized by concentration using the bicinchoninic protein assay (Pierce, 23225). Equal masses of protein were separated, transferred, and analyzed using standard Western Blot techniques with antibodies **(Supp. Table 1)**. The loading ranged from 20 to 50 ug of total protein per lane depending on several variables (gel type, size, well format, sample amount, Ab quality/sensitivity, etc.). Most western blots used Biorad Stain-Free gels (item 5678115) with total protein imaged on the gel after running, and membrane after transfer to quantify equal loading and transfer. Others used Nupage 4-12% Bis-Tris gels (items NP0336).

Images were acquired using enhanced chemiluminescence substrate (Azure, Radiance Plus) on a Biorad Versadoc MP5000 and quantified using Biorad QuantityOne Software (Version 4.6.6) with global background subtraction settings or acquired using chemiluminescence and/or Dual-channel fluorescence on an Invitrogen iBright-1500 and quantified using iBright Analysis Software (Version 4.0.0).

### 4.8 Microtubule Fraction Assay

Powdered heart tissues (25 mg) were homogenized in 0.25 mL microtubule stabilization buffer **(Supp. Table 1)** with a bead mill (Qiagen, TissueLyserII). Homogenates were centrifuged at 1,000×g for 5 mins at 37 ℃. The low-speed pellet was reserved, and 0.1 mL of supernatant was centrifuged at 107,000×g for 60 mins at 37 ℃. The high-speed supernatant was reserved as the free-tubulin fraction, while the high-speed pellet was resuspended in 1% SDS, heated at 90℃, sonicated to denature proteins, and reserved as the polymerized tubulin fraction. Equal volumes of fractions were analyzed using standard western blot techniques with antibodies **(Supp. Table 1)**. Dual-channel fluorescent images were acquired on an Invitrogen iBright-1500 and quantified using iBright Analysis Software (Version 4.0.0).

### 4.9 Tissue Immunofluorescence

Mouse hearts were sectioned on a coronal plane and frozen in OCT medium. Cryostat sections (∼5 μm thick) were affixed to glass slides, postfixed in fresh 4% paraformaldehyde, and permeabilized with 0.2% (v/v) TritonX-100. Slides were blocked, washed, and incubated overnight at room temperature in a humidified chamber with antibodies diluted in blocking buffer **(Supp. Table 1)**. After washing, sections were incubated for 90 mins at room temperature with secondary antibodies diluted in blocking buffer. After washing, slides were mounted with a coverslip using Prolong Diamond (ThermoFisher, P36961) and imaged using a Leica confocal microscope (LSM510) with the 60x oil objective.

### 4.10 Cell Isolation, Culture, and Transfection

Neonatal rat cardiomyocytes (NRCM) were isolated from ∼3-day-old pups from Sprague-Dawley rats (Charles River, Stock 001) following standard protocols using the Worthington Neonatal Cardiomyocyte Isolation System (Worthington, # LK003300) followed by a two-step Percoll density gradient,^15^ and cultured in Cardiomyocyte growth media **(Supp. Table 1)**.

NRCMs were transfected using Lipofectamine 2000 (0.5% v/v, final) with plasmid DNA (2 ng/μL, final) or siRNA (25 nM, Dharmacon SmartPool) diluted in Optimem media at final volume of 100 μL/cm^2^. Sorbs2 expression plasmid was generated by RT-PCR amplification of a mouse cardiac-specific Sorbs2 isoform (primers listed in **Supp. Table 1**; schematic shown in **Supp. Figure 2**, “our clone”) and subsequent cloning into a CMV expression vector. The transfected cells were incubated for four hours before transfection media was removed and replaced with growth media. The cells were incubated for ∼48 hours with daily media exchanges before patch-clamp electrophysiology or calcium imaging experiments.

### 4.11 Determination of cytosolic Ca^2+^ transients

Cells were loaded with Fura-2 acetoxymethyl ester (Fura-2AM) by incubating cells with 1 µM Fura-2AM in HBSS for 20 min at room temperature and incubated at 37° for 20 min to esterify the stain. Cells were excited alternatively at 340 and 380 nm. Fluorescence signal intensity was acquired at 510 nm. Real-time shifts in Fura-2AM fluorescence ratio were recorded about 30 sec before adding an agonist using a Nikon Eclipse Ti2 inverted light microscope. Imaging was acquired every 2 sec to measurement. Peak amplitude (R) was calculated by subtracting the baseline fluorescence ratio from the highest fluorescence ratio. The area under the curve (AUC) was determined using GraphPad Prism and normalized by subtracting the AUC at baseline. Summary data represent the average difference in the basal and peak increase in cytoplasmic [Ca^2+^].

### 4.12 Whole-Cell-Patch Clamp of Na^+^ Currents (INa)

All whole-cell recordings were obtained utilizing the Axon Axopatch 200B amplifier and Digidata 1440B data acquisition system (Molecular Devices), as previously described.^16^ Cell capacitance was recorded after adjusting for transients’ post-membrane rupture. pClamp software (version 10.4) was used for data analysis. To calculate INa density, peak current was divided by the membrane capacitance. To test steady-state activation of INa in NRCMs, a 200-ms prepulse to -120 mV was used to eliminate inactivated channels and cells were subjected to a 200-ms test pulse between -80 mV and +15 mV in increments of 5 mV. Patch pipettes of 2-3 MΩ were used and recipes for internal and extracellular solutions are listed in **(Supp. Table 1)**.

### 4.13 Bioinformatics

Data were exported for additional analyses using Microsoft Excel, Graphpad Prism v8.2.1, and R v3.6.1. RNAseq and RIBOseq data shown in **Supp. Figure 2** were acquired from http://shiny.mdc-berlin.de/cardiac-translatome/,17 or remapped from SRA archive PRJNA477855 ^18^ or SRA archive PRJNA484227.^19^ Relevant datasets of RNA expression **(Figure 1a)** were pulled from GEO (Gene Expression Omnibus), the European Genome-Phenome Archive, or published supplementary data tables. Select data were reanalyzed using GEO2R webtool comparing the difference between cardiomyopathy vs control samples (Datasets described in **Supp. Table 2**). GWAS data **(Table 1)** were acquired from these publications/databases.^3, 20–28^

**Table 1.** Unexplored genetic variation in SORBS gene family underlies cardiovascular diseases.

| RSID | Phenotype | P-value | Reference |
| --- | --- | --- | --- |
| SORBS2 |  |  |  |
| rs5018568 | mRNA eQTL | 9e-6 | 3 GTEx |
|  | P-wave terminal force | 5e-4 | 21 PWI GWAS |
|  | DCM | 0.002 | 22 DCM GWAS |
| rs75898208 | Plasma cTnT | 2e-9 | 23 cTnT GWAS |
|  | Ventricular tachycardia | 2e-3 | 24 UK-Biobank |
| rs10009306 | ICD shocks | 3e-4 | 25 GAME GWAS |
|  | Arrhythmia | 0.06 | 24 UK-Biobank |
| rs182253016 | Hypertensive heart disease | 4e-7 | 24 UK-BioBank |
| SORBS1 |  |  |  |
| rs1410059 | PR-interval | 7e-8 | 26 PR GWAS |
|  | Cardiac arrest | 0.04 | 24 UK-Biobank |
| rs3193970 | Sudden cardiac arrest | 1e-4 | 27 SCA GWAS |
|  | Cardiac arrest | 0.03 | 24 UK-Biobank |
| rs943346 | Cardiovascular disease | 6e-8 | 28 FINDOR |
| rs12221125 | Systolic blood pressure | 9e-9 | 28 FINDOR |

**Figure 1.**
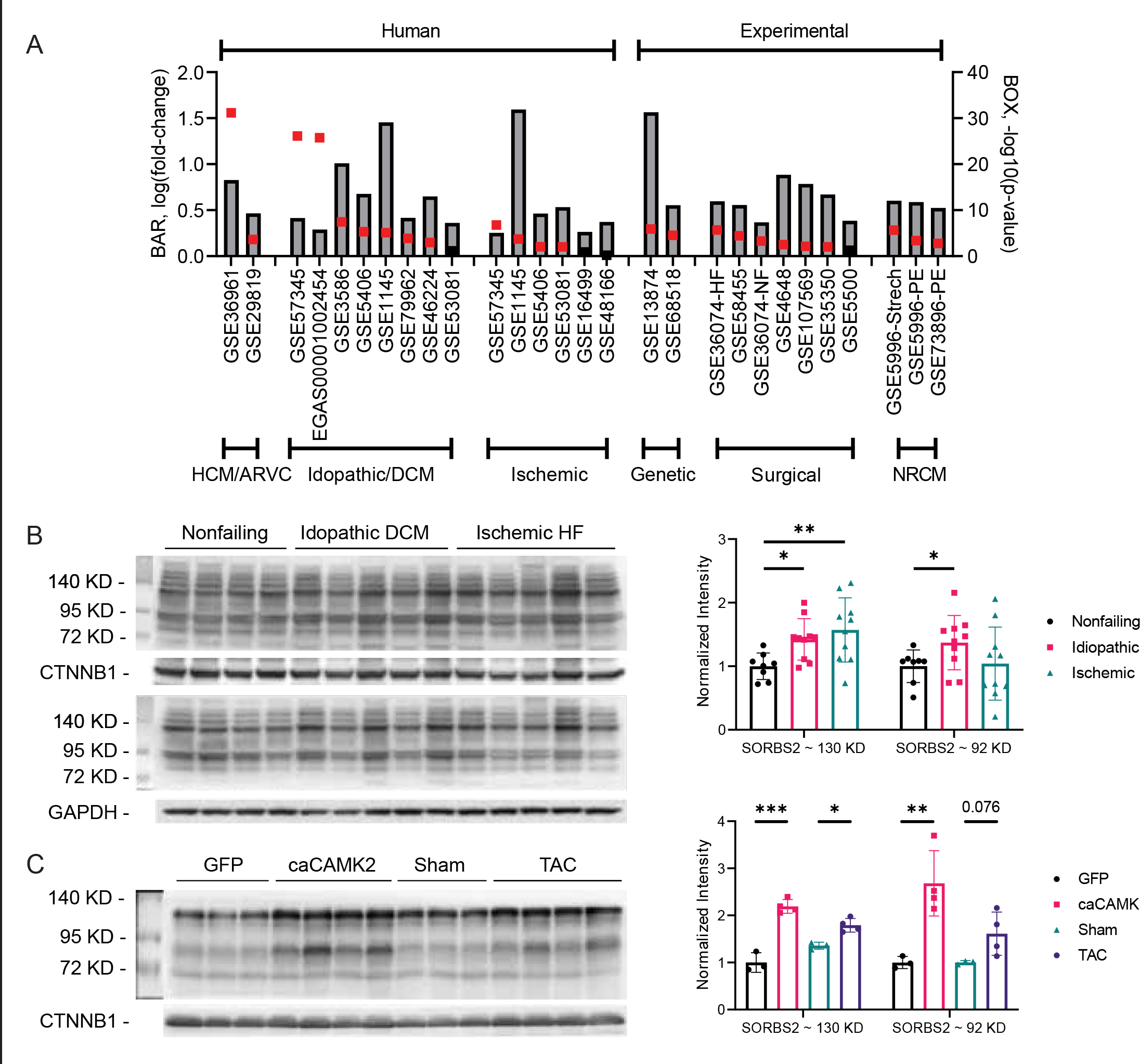
Sorbs2 is broadly dysregulated in adult cardiomyopathies. (A) Plot showing the log2 fold-change and -log10 p-value for Sorbs2 mRNA expression differences between control samples and cardiac disease and/or experimental samples across the indicated human and rodent transcriptional profiling datasets. Red boxes denote p-value < 0.05. Dataset accession numbers and relevant group comparisons are noted, and additional information is in **Supp. Table 2**. (B,C) Representative western blot images and quantitative densitometry analysis for expression of major Sorbs2 protein isoforms (130 and 92 kDa) in human heart samples (B, N=8-10 per group), or mouse cardiac tissues samples collected from mice treated with control AAV-GFP or AAV-caCaMKII, or subjected to either control sham or TAC surgeries (C, N=3-4 per group). Data are plotted as mean +/- SEM. Significance was determined by ANOVA with Dunnett’s post-hoc test, compared to nonfailing controls (B) or by two-tailed t- test between control and disease samples (C), * p<0.05, ** p<0.01, *** p<0.001.

## 5. Results

### 5.1 Sorbs2 is Broadly Dysregulated in Cardiomyopathic Hearts

Previous reports indicate that Sorbs2 expression is increased in heart samples from patients with LVNC,^8^ in serum following myocardial infarction^4^ and genetic mutations in Sorbs2 may underlie ARVC.^7^ However, there are no published efforts interrogating the broad dysregulation of Sorbs2 mRNA expression in diseased hearts. In a systematic analysis of several independent datasets of transcriptome-wide RNA expression, accessed from public data repositories, we examined Sorbs2 expression in non-failing and diseased hearts from human subjects and rodent models. In 13 of 16 datasets from independent human cohort studies, Sorbs2 mRNA expression is significantly increased in patients with ischemic, idiopathic, ARVC, and hypertrophic cardiomyopathy **(Figure 1a)**. In-depth analysis of an available RNA sequencing dataset^18^ revealed that the most abundant cardiac Sorbs2 transcript isoforms are among those significantly increased in dilated and ischemic cardiomyopathy **(Supp. Figure 1a-c)**.

Interestingly, this increase is conserved across species, as Sorbs2 mRNA levels are also consistently higher in 12 of 13 datasets from experimental rodent models of cardiac hypertrophy, ischemia, or genetic cardiomyopathies **(Figure 1a)**.

To assess if Sorbs2 expression is also elevated at the protein level, we performed western blot analyses and found that Sorbs2 protein expression is increased ∼50% in human cardiac tissues from patients with ischemic or idiopathic HF, compared to non-failing control samples **(Figure 1b)**. In mice, Sorbs2 protein expression was also increased ∼2-fold in failing hearts induced by cardiac-targeted overexpression of constitutively-active CaMKII, which elicits severe HF [i.e., reduced ejection fraction (EF) down to ∼20-30%] within three weeks and early death by 7 weeks. In addition, Sorbs2 protein levels were increased by ∼50% in mouse hearts subjected to pressure-overload, resulting in cardiac hypertrophy and mild cardiac dysfunction (i.e., 40-50% increase in heart-weight: body-weight ratio and ∼10-15% decrease in EF) by 8 weeks **(Figure 1c)**.

To assess if Sorbs2 upregulation in HF occurs through transcriptional or posttranscriptional mechanism, we queried ribosomal profiling (RIBOseq) data from human HF^17^. Normalized RNA- and RIBOseq reads were significantly upregulated in nonfailing and DCM samples and calculated translational efficiency (RIBO/RNA ratio) was equal to one independent of disease **(Supp. Figure 1d)**. This suggests that cardiac Sorbs2 transcripts are not likely under substantial translational regulation at baseline or during HF. Altogether, these expression analyses indicate that independent of the etiological origin, myocardial Sorbs2 expression is broadly and consistently increased in the setting of HF, both in human patients and rodent models, likely through transcriptional upregulation or increased transcript stability.

### 5.2 Examination of Sorbs2 Cardiac Isoforms

Sorbs2 is a large gene with complex splicing that gives rise to numerous transcripts [e.g., the Ensembl human genome assembly currently annotates 65 transcripts, including 8 with complete coding sequence (CDS)]. To assess the gene structure of Sorbs2-encoding cardiac mRNA isoforms, we reanalyzed available RNA-seq and RIBOseq data from human and rodent heart tissues^17, 19^ to determine which exons are expressed at the RNA level and translated into protein, focusing on the 8 isoforms harboring complete CDS. Assessment of protein domains across Sorbs2 shows that the characterized cytoskeletal adaptor domains [Sorbin Homology (SoHo) and SH3] are present in each isoform, including those expressed in heart; however, of note, the characterized RNA-binding domain (ZnF-C2H2), is restricted to select transcripts that are not present in cardiac tissue. Along these lines, we examined sequences of Sorbs2 transgenes described in published studies related to cardiomyocyte biology and note that the transgene (Addgene #74514) utilized to demonstrate RNA-binding activity in cardiomyocytes^29^ does not represent a Sorbs2 isoform expressed in cardiomyocytes. The exon harboring the RNA-binding domain is exclusively expressed in neuronal tissues, and indeed, this transgene was originally cloned from mouse brain. While it is clear that much more work is needed to better understand the complexity of Sorbs2 isoforms and their relevance to cardiac biology, this analysis reveals that the cytoskeletal adaptor domains are present in every cardiac isoform, whereas the RNA-binding domain is not expressed in heart.

### 5.3 Generation of Cardiomyocyte-Specific Sorbs2 Knockout Mice

Sorbs2 is broadly expressed in many tissues, with highest expression in cardiac myocyte and smooth muscle containing tissues. To dissect the role of Sorbs2 specifically in cardiomyocytes, we created cardiomyocyte-specific Sorbs2 knockout mice (Sorbs2-cKO) by inter-breeding αMHC-Cre transgenic mice with Sorbs2-fl/fl mice (loxP sites flanking Sorbs2 exon 12) **(Supp. Figure 2b, red box)**. Excision of this obligate exon, present in all Sorbs2 protein-coding transcripts expressed in mouse hearts (TPM>1), will introduce a frameshift and premature termination. Sorbs2-cKO mice are viable and fertile and have a normal development and maturation. Western blot analysis of tissue lysates confirmed the loss of Sorbs2 protein expression in Sorbs2-cKO heart samples, relative to wild-type (WT) controls (Sorbs2-fl/fl, Cre-negative mice), whereas Sorbs2 protein expression persisted in Sorbs2-cKO brains **(Figure 2a)**. Notably, cardiomyocyte loss of Sorbs2 does not lead to a compensatory upregulation of Sorbs1 in heart tissues **(Supp. Figure 3a-b)**. Conditional Sorbs2 deletion was also examined by immunofluorescent staining of cardiac tissue sections, which showed cardiomyocyte-specific loss of Sorbs2 protein in Sorbs2-cKO mice, evidenced by overt lack of Sorbs2-positive staining at the ICD and retained expression in coronary artery smooth muscle **(Figure 2b-c)**. Together, these data support the successful generation of cardiomyocyte-specific Sorbs2 KO mice, with retained expression in other tissues and cardiac cell types.

**Figure 2.**
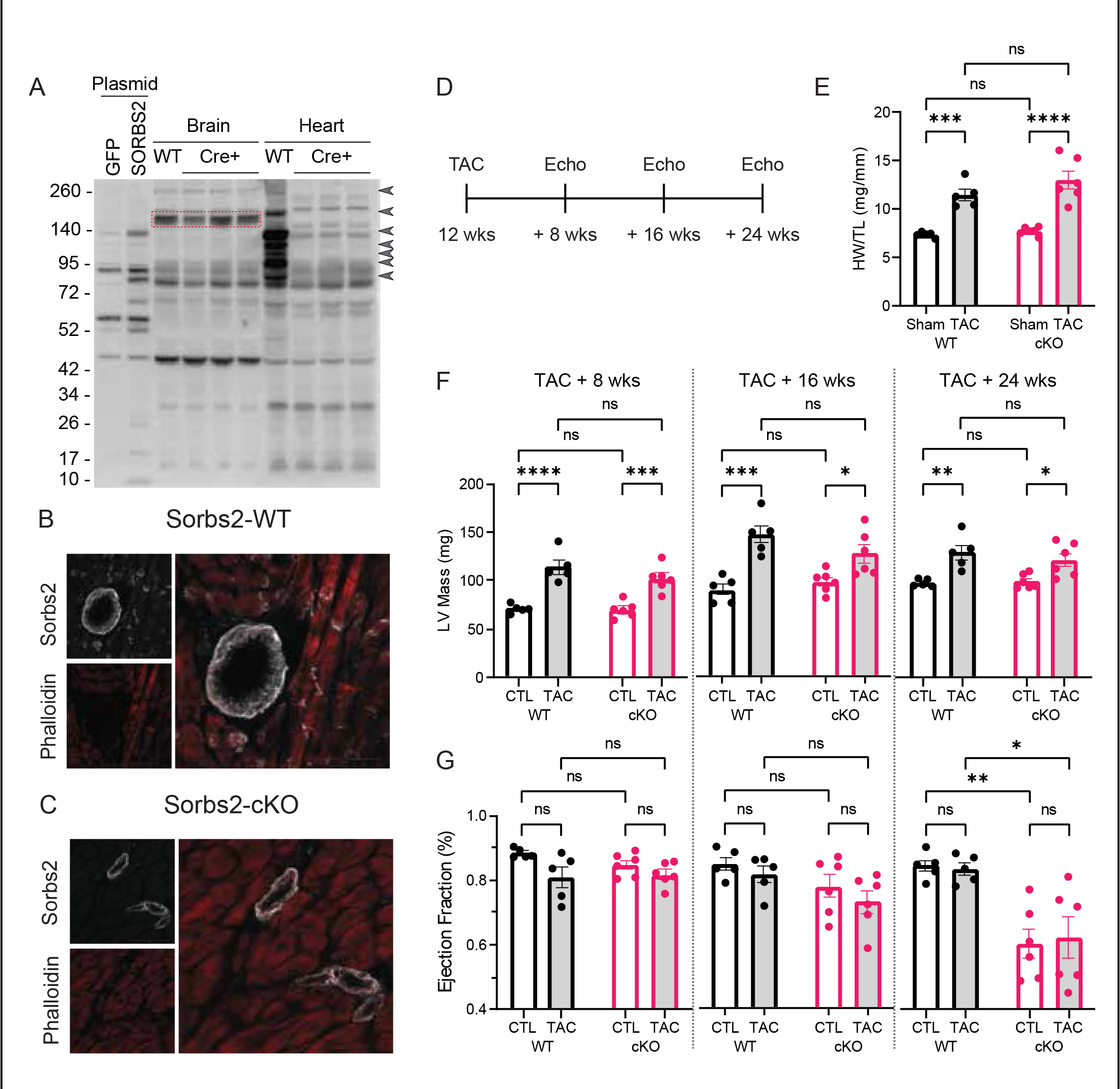
Generation of novel cardiomyocyte-specific Sorbs2 knockout mice. (A) Sorbs2 western blot in brain and heart tissues from wildtype (WT) and Sorbs2 cKO mice. The brain specific Sorbs2 isoform (∼150 kDa, red box) remains expressed in WT and CRE+ samples, whereas several immunoreactive bands are lost in CRE+ heart samples compared to control (arrowheads). The canonical Sorbs2 heart isoform is expected at ∼130 kDa. Lysates from cells transiently-transfected with expression plasmids encoding either Sorbs2 (mouse cardiac- specific isoform, **Supp. Figure 2a**, “our clone”) or GFP as control are also included on the blot for reference. (B-C) Representative Sorbs2 immunofluorescence (white) in heart sections from wildtype (WT) and Sorbs2 cKO mice, co-stained with phalloidin (red). (B) In WT hearts, Sorbs2 is prominently expressed in cardiomyocytes at the intercalated disc (ICD) and in coronary arteries. (C) In cKO hearts, Sorbs2 is expressed in coronary arteries but not in cardiomyocytes. Tissues are from male mice ∼48 weeks old, and scale is 50 μm. (D) Timeline of TAC experiment. (E) Indexed heart mass (HW/TL) at sacrifice in WT and Sorbs2-cKO mice at ∼36 weeks age. (F-G) LV mass and ejection fraction derived from echocardiography at 8-, 16-, and 24-weeks post TAC. Dots show individual mice (N=5-6 group) with mean +/- SEM, statistics acquired using one-way ANOVA with Sidak posthoc test comparing selected groups (each comparison shown on plot), * p<0.05, ** p<0.01, *** p<0.001, **** p<0.0001, ns = not significant.

**Figure 3.**
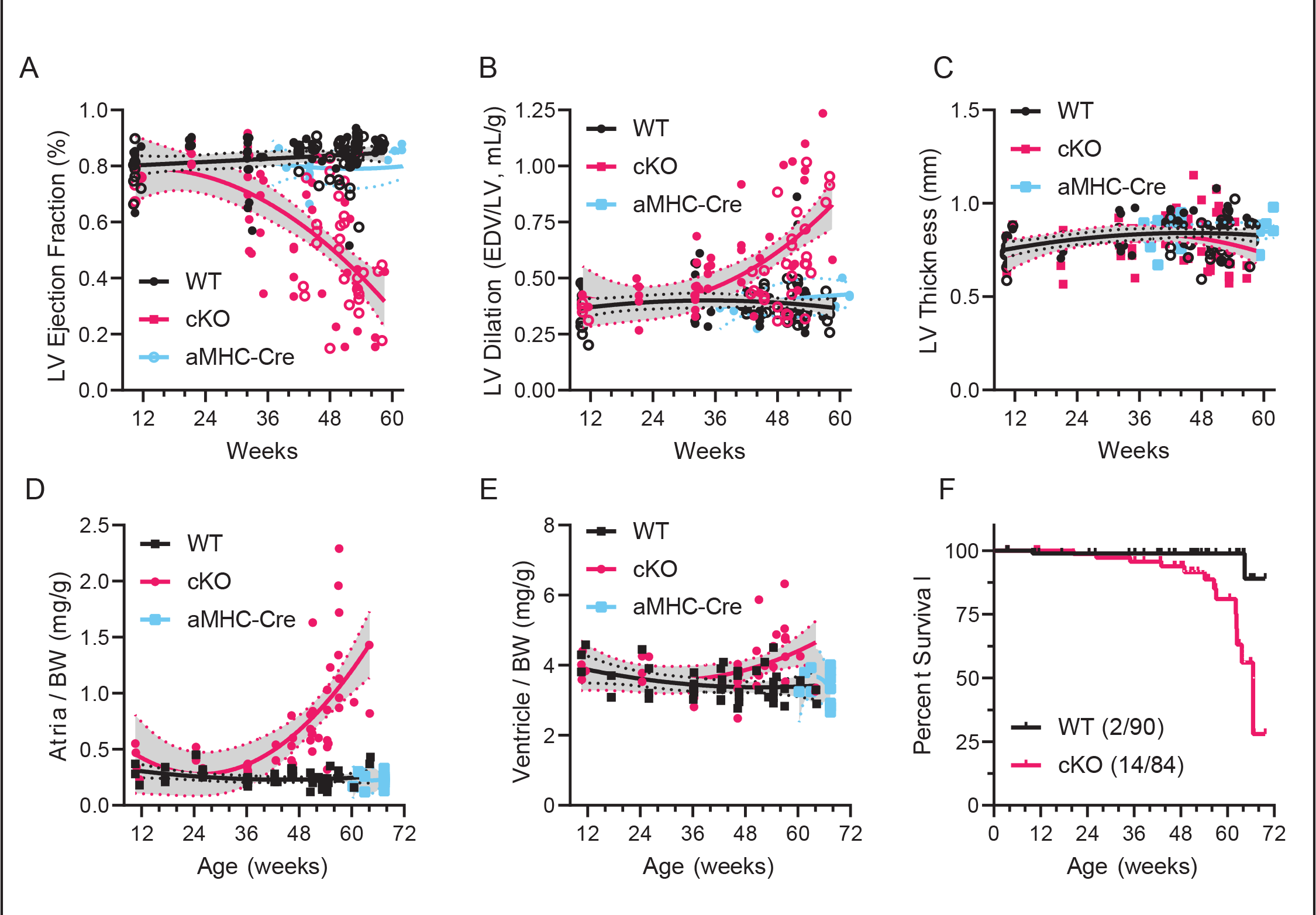
Cardiomyocyte-specific Sorbs2 knockout mice develop age-related systolic dysfunction, cardiac remodeling, and premature death. (A-C) Quantification of echocardiography derived cardiac function over time including (A) LV EF (ejection fraction), (B) indexed LV dilation (LV end diastolic volume divided by LV mass), and (C) LV thickness (long axis view, septum, avoiding papillary muscles); N=74/118 (WT), N=55/93 (cKO), and N=12/48 (αMHC-Cre), N=mice/measurements, some are serial echo on mice over time. Dots represent individual echo/mice (solid=male and open=female) and trendline shows a LOESS nonlinear regression fit across the mix-sexed cohort, with 95% CI shaded grey. (D-E) Post-sacrifice gravimetric analysis of cardiac tissue normalized to body weight including (D) Atria and (E) Ventricle; N=55 (WT), N=46 (cKO), and N=12 (αMHC-Cre), N=male mice. P-values indicate difference between WT and cKO curves analyzed using 2-way ANOVA with Sidak’s multiple comparisons testing the interaction between age and genotype (see **Supp. Table 3)**. (F) Survival curve in mix-sexed cohorts showing premature death of Sorbs2-cKO mice (median survival = 66.57 weeks). Deaths (down steps) represent mice found dead in pen. Censured data (up tics) represent mice euthanized for experiments. Death curves are significantly different using a log-rank test (Mantel-Cox method), N=90 (WT) and 84 (cKO), Chi-square =11.57, and p=7e-4. Total number of deaths are noted in parentheses.

### 5.4 Sorbs2 expression in Cardiomyocytes is not required for cardiac hypertrophy

Given that Sorbs2 is consistently upregulated in HF and cardiac hypertrophy, and that AAV-mediated Sorbs2 overexpression induced cardiac hypertrophy in mice,^8^ we examined the effects of loss of Sorbs2 on the hypertrophic response in male WT and Sorbs2-cKO mice subjected to cardiac pressure-overload by traverse aortic constriction (TAC) at 12 weeks of age **(Figure 2d)**. Serial echocardiography measures (collected at 8, 16, and 24 weeks post-TAC) and gravimetric analyses done at the time of sacrifice (24 weeks post-TAC) showed significant increases in heart size (heart weight normalized to tibia length, HW/TL) and left ventricular (LV) mass in both WT and Sorbs2-cKO mice after TAC, relative to sham surgery controls; however, no genotypic differences were found **(Figure 2e,f)**. Further assessment of echocardiography data to assess cardiac function revealed that, compared to WT mice, both sham control and TAC Sorbs2-cKO mice have significantly decreased ejection fraction (EF) by 24 weeks **(Figure 2g)**. Together, these data suggest that cardiomyocyte Sorbs2 is not required for cardiac hypertrophy and that Sorbs2-cKO mice develop progressive HF independent of pressure-overload.

### 5.5 Cardiomyocyte-Specific Sorbs2 Knockout Mice Develop Age-Related Systolic Dysfunction, Cardiac Remodeling, and Premature Death

To better understand when loss of cardiomyocyte Sorbs2 impairs cardiac function in mice, we performed serial echocardiography in WT (fl/fl-CRE negative), Sorbs2-cKO, and αMHC-Cre only mice from about 10 weeks thru 16 months of age (representative m-mode shown in **Supp. Figure 4a)**. Compared to controls, Sorbs2-cKO hearts show a progressive DCM phenotype characterized by reduced LV EF **(Figure 3a)**, increased LV chamber dilation **(Figure 3b)**, and thinned LV wall thickness **(Figure 3c)**, with significant differences first appearing after 30 weeks in EF, 40 weeks in dilation, and 50 weeks in wall thickness **(Supp. Table 3)**. Sorbs2- cKO male mice develop worse systolic dysfunction, earlier compared to female mice. Left atrial (LA) enlargement becomes evident on echo past 10 months of age and several mice at a year of age presented with a large LA thrombus, likely arising from blood stasis due to decreased atrial function, considering no evidence for atrial fibrillation, mitral regurgitation, or valve stenosis in these mice. Sorbs2-cKO mice trended toward increased RV thickness **(Supp. Figure 4b)** coincident with LV dysfunction; however, RV dysfunction, RV chamber dilation, or RA enlargement were not evident in echocardiographic analyses until end-stage congestive HF, in contrast to Sorbs2 global KO mice. Gross dissection revealed that Sorbs2-cKO mice have enlarged hearts with dilated LA and LV. Four-chamber gravimetric analyses of WT and Sorbs2- cKO hearts sacrificed at 48 weeks of age indicate that loss of Sorbs2 causes selective significant increases in LA mass **(Supp. Figure 4c)**. Time-course analysis of atrial size shows that Sorbs2- cKO mice have significantly increased total atrial size (total atria mass normalized to body weight) as early as 3-5 months of age **(Supp. Figure 4d)**, with substantial increases after 30 weeks of age **(Figure 3d)**. This increase is primarily driven by LA mass versus RA or bi-atrial masses (cKO regression, r^2^=0.596, p=2e^-4^) **(Supp. Figure 4e)**. Total ventricular mass is significantly correlated with the increased atrial mass (cKO regression, r^2^=0.479, p<1e^-4^) **(Supp. Figure 4f)**; however, the magnitude of ventricular hypertrophy is substantially less **(Figure 3e)**, reaching significance after 50 weeks **(Supp. Table 3)**. Notably, transgenic αMHC-Cre only control mice did not develop large atria or ventricles, or signs of HF **(Figure 3)**^30^. Coincident with worsening cardiac structure and function, Sorbs2-cKO mice exhibit shortened lifespans with premature death starting at ∼one year of age **(Figure 3f),** (males at ∼12 months, females at ∼14-16 months). Both sexes ultimately succumb after ventricular pump function fails and mice develop pericardial and pleural effusions and ascites, indicative of congestive HF. Altogether, these data indicate that Sorbs2-cKO mice develop progressive DCM phenotype with systolic dysfunction, cardiac remodeling, and premature death.

**Figure 4.**
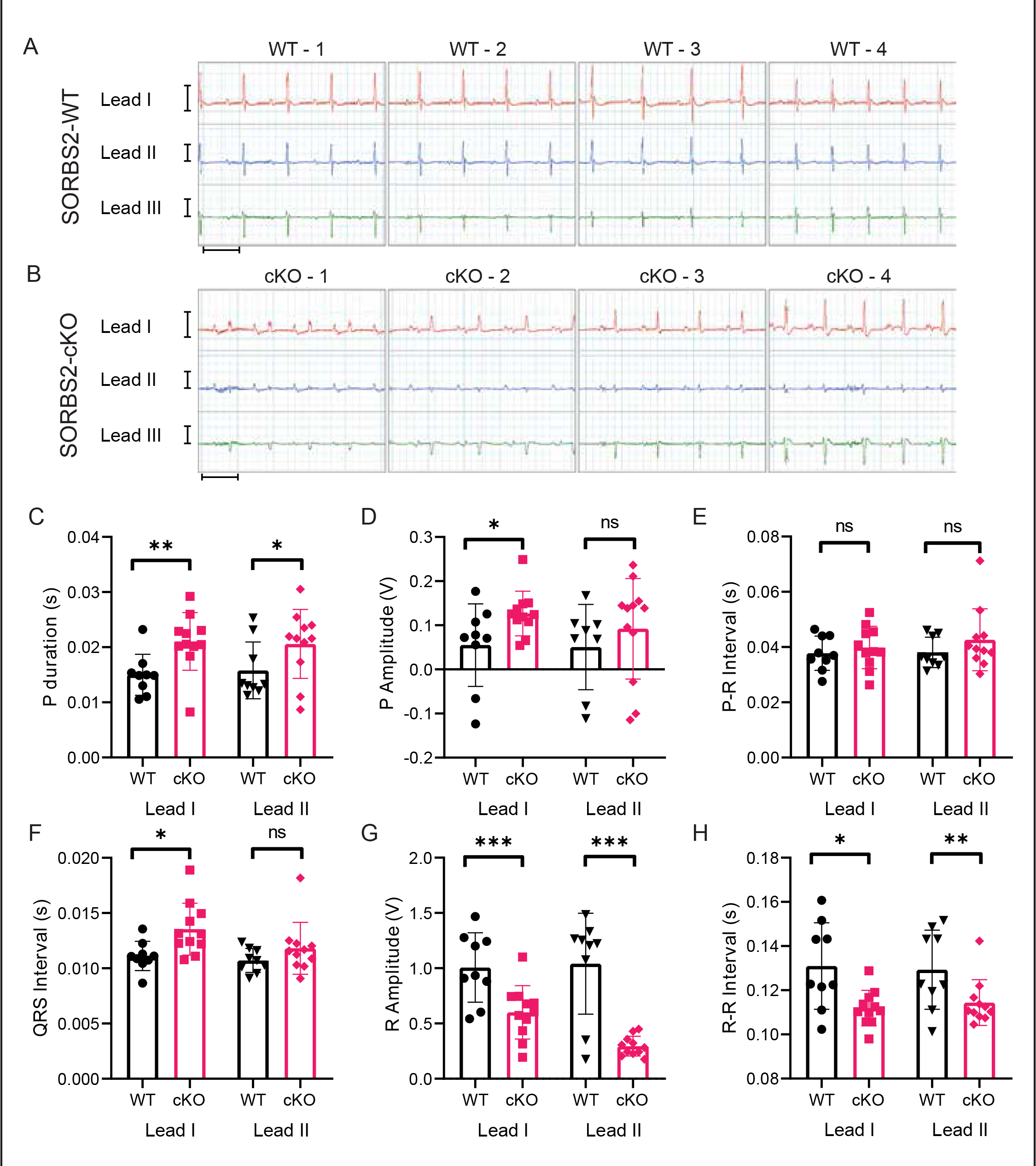
Cardiomyocyte-specific Sorbs2 knockout mice display both atrial and ventricle conduction deficiencies without A-V block. (A-B) Representative multi-lead surface EKG recordings from (A) Sorbs2-WT and (B) Sorbs2- cKO mice aged ∼12 months old. Sorbs2-cKO mice maintain sinus rhythm but show obvious bifid P-waves, increased P-wave and QRS duration, and decreased R-wave amplitude. Y-scale = 1 V, X-scale = 100 ms. (C-H) Quantification of the indicated EKG parameters from WT and cKO mice aged ∼3 months old. Dots represent individual mice with mean +/- SEM, N=9 (WT) and N=11 (cKO), significance from T-Test comparing WT and cKO for Lead-I or Lead-II, * p<0.05, ** p<0.01, *** p<0.001, ns=not significant.

### 5.6 Cardiomyocyte-Specific Sorbs2 Knockout Mice Have Abnormal Cardiac Electrophysiology without A-V Block

Given the prominent localization of Sorbs2 at ICDs, we hypothesized that loss of cardiomyocyte Sorbs2 may disrupt cardiac electrophysiology via initiation or conduction of electrical signals through the heart. We recorded surface EKGs in WT control and Sorbs2-cKO mice from 3 months through one year of age **(Figure 4a-b)**. Collectively, these data reveal that loss of Sorbs2 in cardiomyocytes causes an obvious bifid P-wave morphology in all three leads (**Figure 4b)** with increases in P-wave duration **(Figure 4c)** and P-wave amplitude **(Figure 4d)**, starting at 3 months and worsening with age. There was no change, however, in PR interval **(Figure 4e)**. In Sorbs2-cKO hearts, QRS duration is increased **(Figure 4f)** and R amplitude is reduced **(Figure 4g)**. Heart rates did subtly increase in Sorbs2-cKO mice (derived from R-R intervals, **Figure 4h)**, perhaps to compensate cardiac output. Overall, the EKG morphologies in Sorbs2-cKO mice suggest slow conduction in the atria (increased P-duration) and slow conduction in the ventricles (increased QRS, decreased R-amp); however, major conduction defects (i.e., A-V block) or significant tachy- and brady- arrythmias including atrial fibrillation and ventricular tachycardia were not found. Instead, the slow atrial conduction and abnormal p-wave morphology may reflect left atrial enlargement **(Supp. Figure 4c-f)**, a common cause of bifid p-waves in clinical practice (i.e., P mitrale).^31^ Together, these data show that Sorbs2-cKO mice may have abnormal electrophysiology within the atria and ventricles, perhaps due to structural remodeling, without showing significant atria-to-ventricle conduction deficiencies (i.e., A-V block).

### 5.7 Cardiomyocyte-Specific Sorbs2 Knockout Mice Exhibit Age-Related Reductions in Cardiac and Myofiber Contractility

Sorbs2 is localized within cardiomyocytes at intercalated disks, Z-disks, and costameres, where it primarily crosslinks cytoskeletal components and associated signaling complexes. Our data show that cardiomyocyte-specific loss of Sorbs2 in mice leads to a clear DCM phenotype by one year of age. Given that loss of Sorbs2 may weaken the cardiomyocyte cytoskeletal architecture, we hypothesized that Sorbs2-cKO mice hearts would have reduced contractility, prior to detectable systolic dysfunction on echocardiography. To test this, we catheterized ∼25- week-old mice and measured LV hemodynamics at baseline and during a dobutamine (DOB) infusion (dose range 2-12 ng/g/min). WT control mice (fl/fl, Cre negative) increased cardiac contractility (shown as dP/dTmax) coincident with increasing DOB concentrations, whereas Sorbs2-cKO mice did not **(Figure 5a)**. Both genotypes similarly increased heart rate in response to DOB challenge **(Supp. Figure 5a)**, indicating that adrenergic signaling was intact in Sorbs2- cKO hearts. At baseline, cardiac contractility was not different among mice; however, control mice achieved a sustained increase in dP/dTmax, with peak contractility ∼1400 mmHg/sec at 12 ng/g/min DOB, the highest dose tested **(Figure 5b)**. Contractility was significantly reduced in Sorbs2-cKO mice with a peak at 8 ng/g/min and remained blunted through 12 ng/g/min to ∼9000 mmHg/sec **(Figure 5b)**. Cardiac relaxation (indicated by dP/dTmin) was not different between groups at baseline but was significantly blunted in Sorbs2-cKO mice during DOB challenge **(Supp. Figure 5b)**. In addition, at baseline, Sorbs2-cKO hearts showed increased end-diastolic pressure (EDP), which further increased with DOB **(Figure 5c)**, suggesting basal diastolic dysfunction. Surface EKG measurements were also recorded from both WT control and Sorbs2- cKO mice during the DOB challenge. Although bifid P-waves and increased P-durations were present in Sorbs2-cKO mice, they maintained normal sinus rhythm with no signs of atrial or ventricular arrythmias or A-V block **(Supp. Figure 5c-f)**.

**Figure 5.**
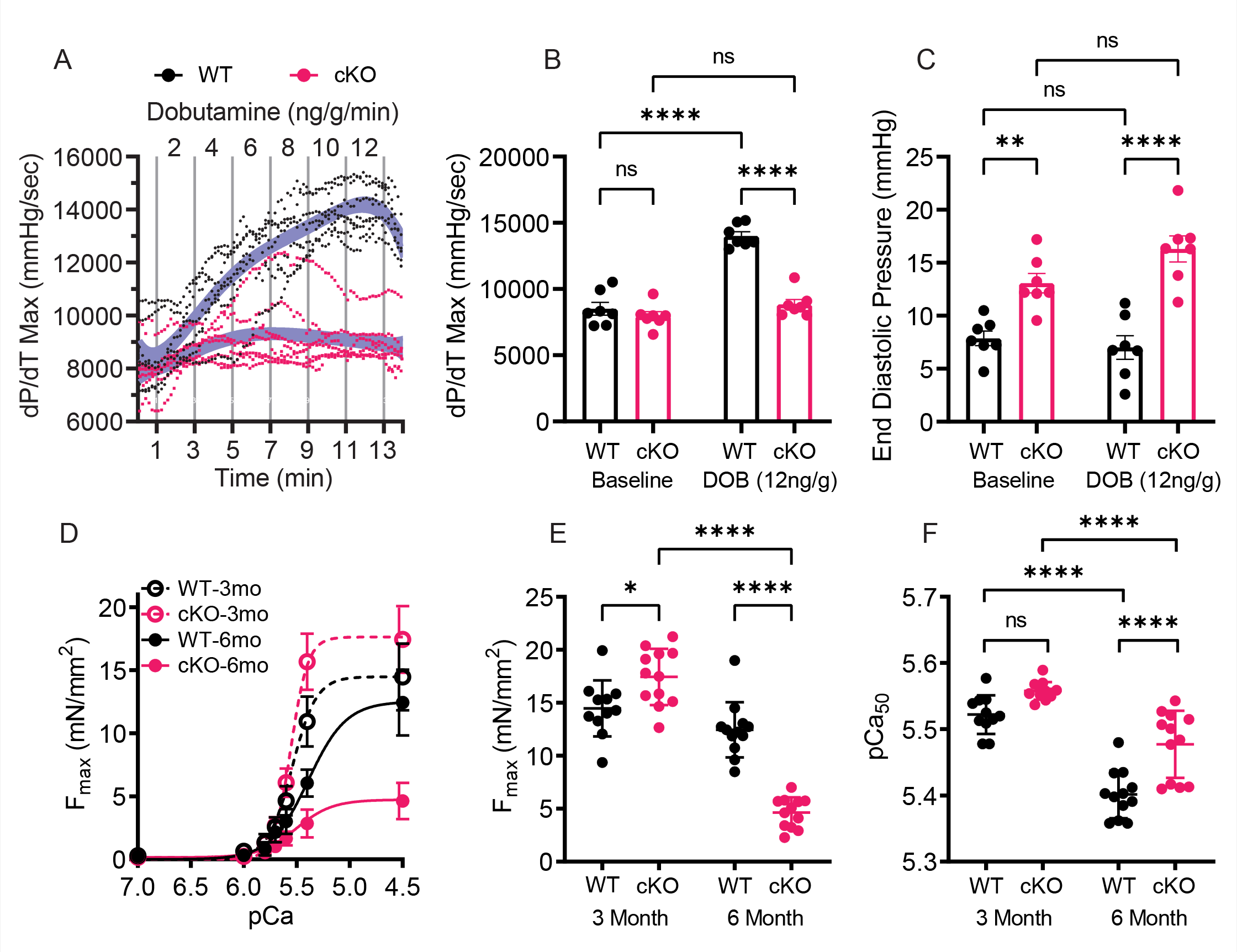
Cardiomyocyte-specific Sorbs2 knockout mice have severe contractile dysfunction. (A) Cardiac contractility (dP/dT Max) derived from LV catheterization and stepped infusion of dobutamine (DOB, 2 min per dose indicated on top) in ∼6-month male mice. Dots show 10 second average for individual mice (n=7 mice per group). The blue shading denotes a LOESS nonlinear regression +/- 95% confidence interval and are significantly different (p<1e-4) via repeated measures 2-way ANOVA with Sidak’s multiple comparisons testing the interaction between age and genotype. Final 30 second average per DOB dose (12 ng/g/min) for (B) dP/dT Max (C) and end diastolic pressure (EDP). (F) Isometric pCa-tension curves generated from skinned LV myofibers male mice at 3 and 6 months old. Each datapoint represents the mean +/- SEM of 11-12 myofibers isolated from 4 different mice per group (2-3 fibers per mouse). Trendline denotes a least-squares fit to sigmoidal dose-response curve with variable slope. (G) Maximum developed tension (Fmax, mN/mm2) (H) Calcium sensitivity (pCa50, M) were calculated from isometric pCa-Tension curves. (B, C, E, F) Dots show individual data (mice or myofibers) with mean +/- SEM, statistics acquired using one-way ANOVA with Sidak posthoc test comparing selected groups (each comparison shown on plot), * p<0.05, ** p<0.01, *** p<0.001, **** p<0.0001, ns = not significant.

DOB challenge data suggest that Sorbs2-cKO hearts have intrinsic defects in contractility causing insufficient contraction within cardiomyocytes. To test this, calcium-induced isometric tension was measured in permeabilized LV myofibers from WT (fl/fl, CRE negative) and Sorbs2-cKO mouse hearts at 3 or 6 months of age **(Figure 5d)**. Contrary to our hypothesis, these data show that myofibers from 3-month-old Sorbs2-cKO hearts do not have reduced maximal force generation, nor reduced calcium sensitivity, but instead show increased maximum tension **(Figure 5e-f)**, perhaps due to compensatory mechanisms. In contrast, LV myofibers from 6- month-old Sorbs2-cKO mice show a significant reduction in tension development, with minor differences in calcium sensitivity **(Figure 5e-f)**. These findings indicate that prior to systolic dysfunction, Sorbs2-cKO mice have impaired myofiber contractility, consistent with our DOB challenge findings. Altogether, the data support that appropriate cardiomyocyte excitation remains intact during beta-adrenergic stress in Sorbs2-cKO mice; however, these mice fail to increase cardiomyocyte mechanics and/or couple excitation to contraction.

### 5.8 Molecular Changes in Cardiomyocyte-Specific Sorbs2 Knockout Hearts

Sorbs2-cKO mice develop age-dependent DCM with abnormal electrophysiology and contractility, however the underlying molecular mechanisms remain undefined. We explored several possibilities to determine how loss of Sorbs2 over time causes HF in mice, focusing on ICDs, ion channels, calcium handling, and cytoskeletal/sarcomeric proteins. Relevant summarized time-course western blot data (done on WT and Sorbs2-cKO heart lysates collected at 3, 6 and 12 months of age) are shown in **Figure 6a-d**, and representative data shown in **Supp. Figure 6a-d**.

**Figure 6.**
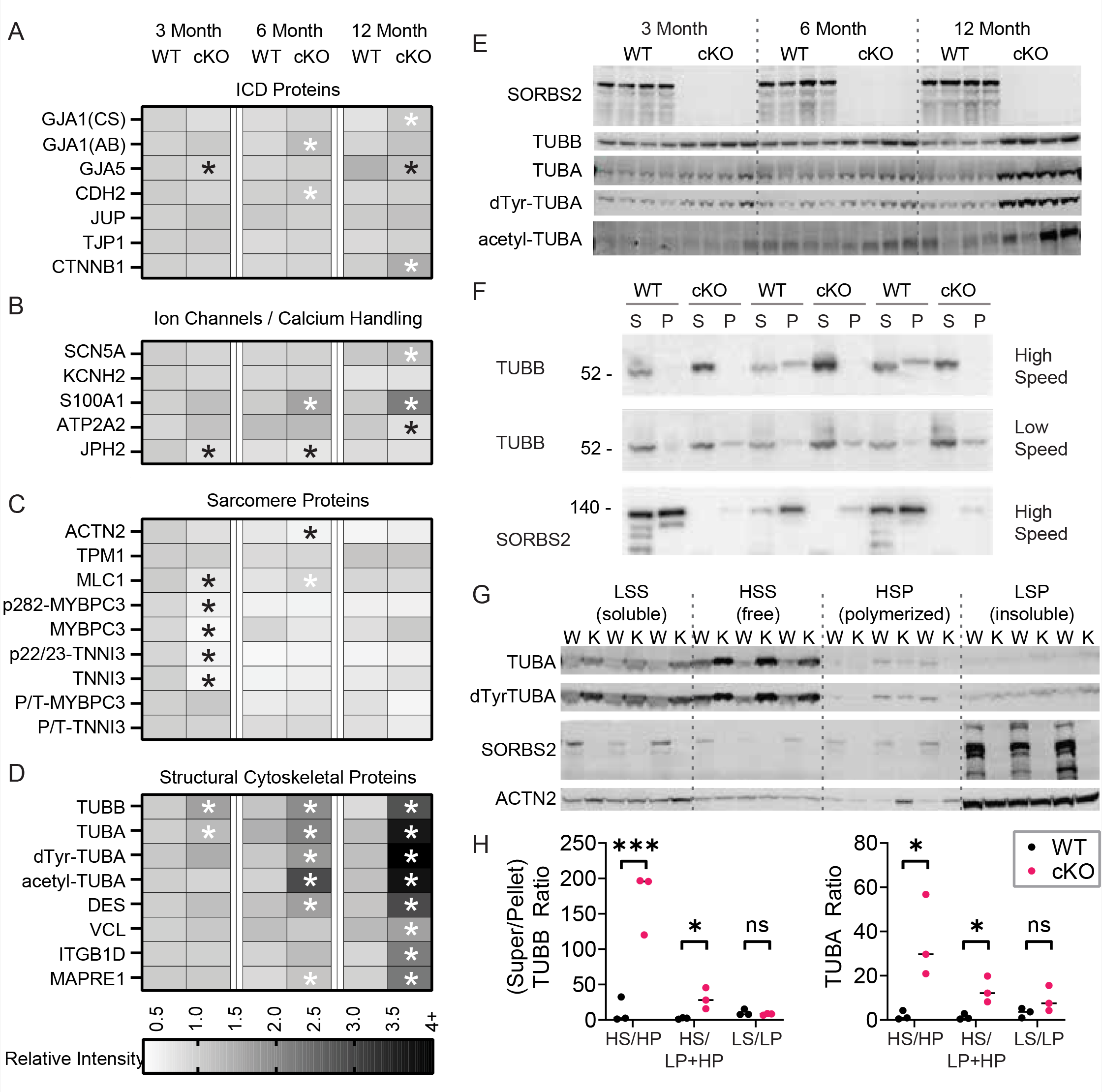
Cardiomyocyte-specific Sorbs2 knockout mice have dysregulated cytoskeletal protein expression not ICD. (A-D) Heatmaps show analysis of protein expression from WT and Sorbs2-cKO cardiac lysates at 3, 6, and 12 months of age. Each box represents the mean integrated intensity, normalized to loading control and expressed relative to 3-month WT; n=4 mice per group. Significant differences (p<0.05) denoted with (*, black for down-regulated and white for up-regulated) overlay on the heatmap, were acquired using t-test comparing cKO to WT at each age. (Abbreviations: CS = Cell Signaling, AB = Abclonal). Raw western blots are shown in **Supp. Figure 6**. Heatmaps are organized by (A) intercalated disc (ICD) proteins, (B) ion channels and calcium handling proteins, (C) sarcomere proteins, and (D) structural cytoskeletal proteins. (E) Western blots show Sorbs2-cKO hearts have significant, early, and sustained expression of microtubule proteins (quantified data in panel 6D). (F) Microtubule fractionation assay from cardiac tissue for male mice aged 16 weeks shows decreased level of polymerized Tubb and increased levels of free Tubb high speed fractions from Sorbs2-cKO hearts. Sorbs2 is shown to be present in high speed, polymerized microtubule fraction. (G) Microtubule fractionation assay also shows decreased level of polymerized microtubule proteins (Tuba, dTyr-Tuba) and increased levels of free microtubule proteins high-speed fractions from Sorbs2-cKO hearts. Note the majority of Sorbs2 is retained in the low-speed pellet, representing 1% Triton-X insoluble proteins, along with sarcomeric actinin and likely other cytoskeletal proteins. (H) Ratiometric quantitation of free/polymerized Tubb and Tuba. Dots represent individual mice with mean, statistics acquired using T-test comparing cKO to WT, * p<0.05, *** p<0.001, ns = not significant. (Abbreviations: S=supernatant, P=pellet, H=high speed, L=low speed)

#### 5.8.1 Cardiomyocyte-Specific Sorbs2 Knockout Retain ICD Protein Expression and Localization

Prior work suggests that Sorbs2 is an RNA-binding protein that is required to stabilize Gja1 (Connexin 43, Cx43) mRNA and maintain ICD structure.^7, 29^ To assess if Sorbs2-cKO hearts have less Gja1/Cx43 or altered expression of other ICD proteins, we measured protein levels via western blot **(Figure 6a, Supp. Figure 6a)**. Despite prior observations that global Sorbs2 KO hearts show a >90% reduction in cardiac Cx43 protein levels,^7^ surprisingly, Sorbs2- cKO hearts maintain normal Cx43 expression from 3 to 12 months of age. In addition, no other ICD proteins, including Connexin 40 (Gja5), N-cadherin (Cdh2), gamma-catenin (Jup), tight junction protein 1 (Tjp1/Zo1), and beta catenin (Ctnnb1), were found to be consistently altered. Overall, these data support that loss of Sorbs2 in cardiac myocytes does not grossly perturb expression of key ICD proteins and further contradicts (along with RNA-seq our analyses described above) the notion that Sorbs2 is an RNA-binding protein that promotes expression of Gja1/Cx43 or other ICD proteins in cardiomyocytes.

Cardiac conduction and contractile deficits are linked to mutations in cytoskeletal and ICD genes, and derangements in cardiomyocyte cytoskeletal architecture. Global Sorbs2 KO mice show overt downregulation of Gja1/Cx43 and mis-localization of other ICD proteins.^7, 29^ To test if cardiomyocyte-specific loss of Sorbs2 causes similar aberrations, we stained cytoskeletal proteins in longitudinal heart sections from WT control and Sorbs2-cKO mice at 8 months of age **(Supp. Figure 7a-b)**. Cell membranes stained with wheat germ agglutinin (WGA) show myocytes of consistent size and shape in both genotypes. WT hearts show Sorbs2 colocalized with beta-catenin (Ctnnb1) at ICD structures and in striations along lateral membranes consistent with costamere structures **(Supp. Figure 7a)**. Sorbs2-cKO hearts lose immunoreactivity for Sorbs2 at ICDs and costameres, however cardiomyocytes maintain normal Ctnnb1 localization at ICDs **(Supp. Figure 7b)**. Staining of heart sections with Phalloidin, and antibodies against sarcomeric actinin, vinculin (Vcl), and Gja1/Cx43 also showed consistent cardiomyocyte size and sarcomere distribution, and remarkably, Vcl and Gja1/Cx43 expression and localization at costameres and/or ICDs were no different between control and Sorbs2-cKO hearts **(Supp. Figure 7c-f)**.

**Figure 7.**
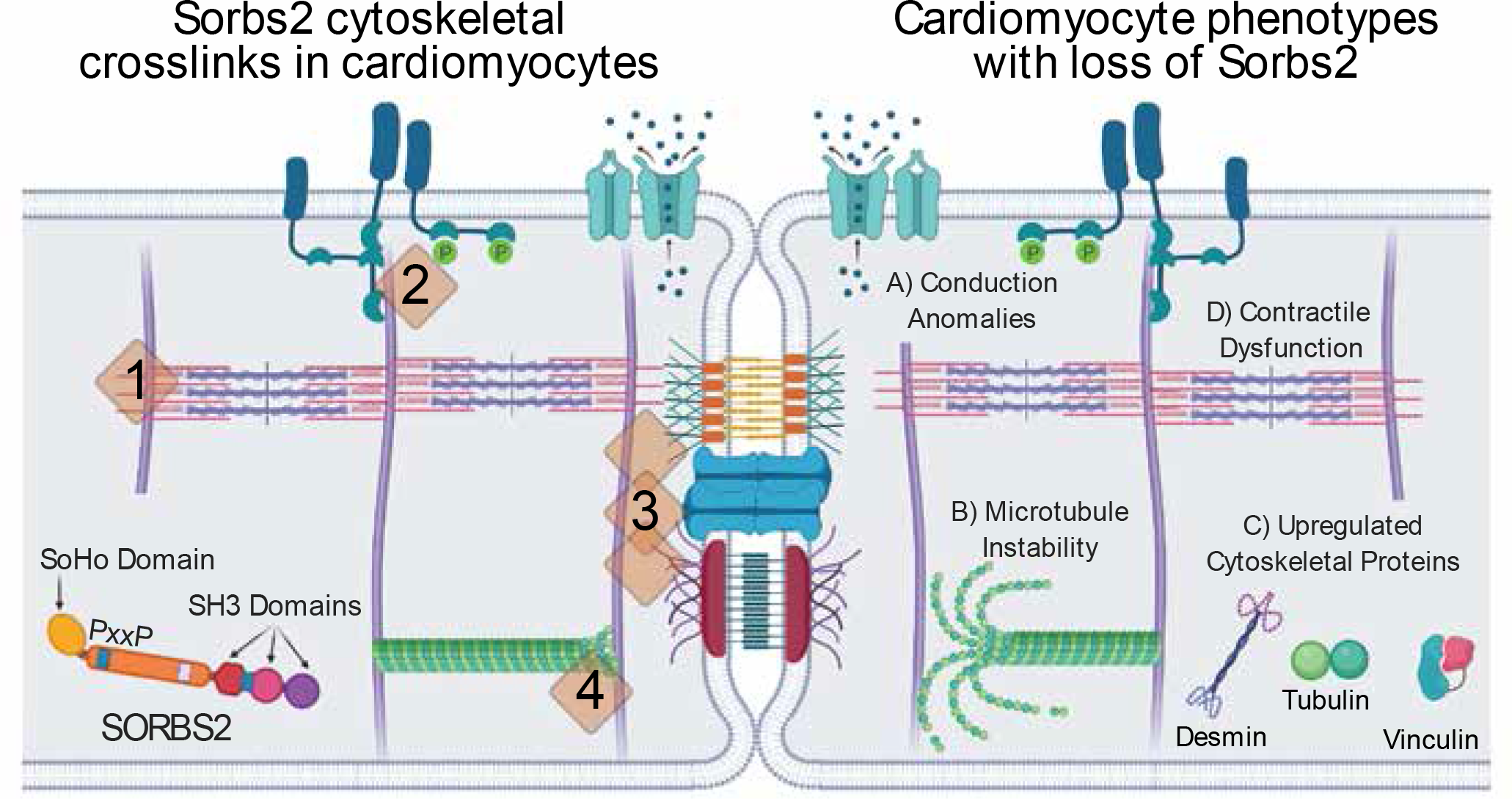
Theoretical model of Sorbs2 functional interactions in cardiomyocytes. Sorbs2 is a cytoskeletal adaptor protein that facilitates diverse protein-protein interactions through its SoHo, SH3 domains, and proline rich motifs (PxxP). Sorbs2 functional interactions in cardiomyocytes include 1) z-discs, 2) costameres, 3) intercalated discs, and 4) microtubules, through crosslink binding with intermediate filaments, actinin and other adaptor proteins, and cell signaling complexes. Age-dependent dilated cardiomyopathy occurs after cardiomyocyte-specific deletion of Sorbs2, which broadly weakens these cytoskeleton structures, and manifests as early A) conduction anomalies, in part, caused by atrial enlargement, B) microtubule instability, C) increased cytoskeletal protein expression, and eventually D) contractile dysfunction leading to HF. Created with BioRender.com.

#### 5.8.2 Ion Channels and Calcium Handling

##### Ion Channels

Sorbs2-cKO mice have abnormal cardiac electrophysiology with increased p-wave duration and amplitude, which may be due to an atrial conduction deficiency, abnormal depolarization, or increased atrial size. The voltage-gated sodium channel Nav1.5 (encoded by Scn5a) is a primary contributor to cardiac depolarization,^32^ Sorbs2 is known to cluster membrane associated complexes^33, 34^ and predicted to bind Nav1.5, and co-expression analyses strongly support that Sorbs2 and Scn5a expressions are highly correlated. Thus, we tested whether loss of Sorbs2 decreases Scn5a expression or regulates Nav1.5 activity.

Unexpectedly, western blot analysis shows that 12-month-old Sorbs2-cKO hearts trend toward increased Nav1.5 expression compared to WT mice, coinciding with the DCM phenotype, and end-stage HF **(Figure 6b, Supp. Figure 6b)**. Expression of Kcnh2 (encodes the cardiac inward rectifying potassium channel mERG, which is primarily responsible for cardiac repolarization) is also unchanged in Sorbs2-cKO hearts. Scn5a and Kcnh2 encode two of several ion channels downregulated in global Sorbs2 KO mice and purported to be direct targets for Sorbs2 RNA-binding activity;^29^ however, our data contradicts this conclusion.

Next, electrophysiology experiments were done using neonatal rat cardiomyocytes (NRCMs) transiently-transfected with expression plasmids encoding either Sorbs2 (mouse cardiac-specific isoform, **Supp. Figure 2a**, “our clone”) or GFP as control and assessed for endogenous sodium current (INa). In parallel, NRCMs were transiently-transfected with non-targeting control or Sorbs2 siRNAs for loss-of-function comparisons. The resulting data show that Nav1.5 channel current density is not affected by overexpression or knockdown of Sorbs2 **(Supp. Figure 8c-f**). Together, these data suggest that potential Sorbs2-mediated regulation of Nav1.5 expression or activity likely does not account for the conduction deficits observed in Sorbs2-cKO mice.

###### Calcium Handling

Sorbs2-cKO mice have abnormal contractility that may be due to dysregulated calcium handling/signaling in cardiomyocytes. We assessed the potential for Sorbs2-cKO hearts to exhibit decreased expressions of S100a1 (multifactorial calcium binding protein), Atp2a2 (Serca2a, calcium transporter), and Jph2 (junctophilin 2, cardiac dyad scaffolding protein). Jph2 levels were only slightly decreased at 3 and 6 month, Atp2a2 expression is decreased at 12 months **(Figure 6b, Supp. Figure 6b)**, and no changes in the ratio of phosphorylated/total Tnni3 and Mybpc3, a hallmark of calcium-dependent posttranslational regulation, were found in Sorbs2-cKO hearts **(Figure 6c, Supp. Figure 6c)**. Unexpectedly, we found that myocardial S100a1 expression is increased 2-3 fold in Sorbs2-cKO mice at 6 and 12 months of age (trending at 3 months); this is interesting and perhaps another indication of compensation, considering that S100a1 bolsters cardiac function in the setting of HF.^35, 36^

To begin addressing whether decreased Sorbs2 expression in cardiomyocytes alters calcium, we transiently transfected NRCMs with siRNA against Sorbs2 and measured whole-cell basal calcium. Within 48 h, Sorbs2 knockdown led to significant increases in whole-cell calcium concentrations **(Supp. Figure 8b**), which compliments published findings showing that acute overexpression of Sorbs2 decreases peak intracellular calcium; however, it is important to note that the latter was deemed to occur secondary to Sorbs2-mediated densification of microtubule networks.^8^

##### 5.8.3 Sarcomere Proteins

Next, we assessed if loss of cardiomyocyte Sorbs2 alters expression or phosphorylation of sarcomeric and related calcium-regulated proteins **(Figure 6c, Supp. Figure 6c)**.

Interestingly, these data show that Sorbs2-cKO hearts exhibit an early (at 3 mo.) decrease in levels of several sarcomere-related protein (myosin light chain, troponin I, or myosin binding protein c3) which is mostly “normalized” by 6 months of age, and not different at 12 months of age. In addition, the calcium-dependent phosphorylation status (i.e., ratio of phosphorylated/total) of p22/23-Tnni3 and p282-Mybpc3 is unchanged between WT and Sorbs2- cKO hearts at 3, 6, or 12 months of age. Staining of actin fibers in heart sections with Phalloidin and antibodies against sarcomeric actinin, showed consistent cardiomyocyte size and sarcomere distribution between control and Sorbs2-cKO hearts **(Supp. Figure 7c-d)**. Together, these data support that sarcomeric derangements and calcium sensitivities are not likely culprit triggers for cardiac dysfunction in Sorbs2-cKO mice.

##### 5.8.4 Sorbs2-cKO mice exhibit derangements in cardiac structural cytoskeletal protein expression and destabilized microtubules

Beyond the potential for ICD derangements, recent reports indicate that Sorbs2 binds cytoskeletal proteins, including tubulins, and regulates microtubule dynamics in cardiomyocytes.^8, 37^ Indeed, across all of the proteins/pathways that we assayed by western blot, the most profound, earliest, and consistent changes occurred with microtubule proteins, with Sorbs2-cKO hearts showing significantly increased levels of beta-tubulin (Tubb), alpha-tubulin (Tuba) detyrosinated-Tuba and acetylated-Tuba **(Figure 6d-e)** at 3 months of age and persisting through 6 and 12 months. The microtubule adaptor protein EB1 (Mapre1) and integrin protein Itgb1d, which modulate microtubules dynamics, increased in similar fashion. Other noncontractile cytoskeletal proteins including Des and Vcl, were also elevated after 6 or 12 months, respectively **(Figure 6d, Supp. Figure 6d)**; however, these are common downstream hallmarks of contractile dysfunction found in most cardiomyopathies.

To determine if cardiomyocyte-specific loss of Sorbs2 alters microtubule polymerization/stabilization, we performed western blot analyses after subcellular fractionation of WT and Sorbs2-cKO heart lysates from ∼16-week-old mice. This revealed that Sorbs2-cKO samples have significantly increased free Tubb and decreased polymerized Tubb **(Figure 6f, h)**. Tuba and detyrosinated Tuba also show increased free/polymerized ratio in Sorbs2-cKO samples **(Figure 6g-h),** suggesting defective microtubule polymerization or stabilization. While the vast majority of Sorbs2 in these samples remained in the low-speed pellet, likely representing detergent insoluble proteins, including other cytoskeletal components (see sarcomere actinin for comparison), some Sorbs2 was notably found in high-speed pellet fractions **(Figure 6f-g)**, consistent with Sorbs2 attachment to polymerized microtubules. Altogether, these data provide a loss-of-function complement to published findings showing that Sorbs2 overexpression promotes microtubule polymerization/stability and suggest that early and persistent changes in microtubule proteins could be at the root of cardiac dysfunction in Sorbs2-cKO mice.

##### 5.8.5 Summary of Molecular Changes in Sorbs2-cKO Hearts

Overall, our investigations into potential underlying mechanisms do not indicate significant redistribution of ICD proteins, rearrangement of sarcomeres, or dysregulated expression of ICD proteins in mice with cardiac-specific loss of Sorbs2. This is consistent with others who show that while Sorbs2 is a component of tight and adherens junctions in epithelial cells, Sorbs2 loss does not affect the assembly, structure or function of these junctions.^38^ Rather, our findings support that decreasing heart function in Sorbs2-cKO mice is preceded by early abnormalities in microtubule dynamics, which associates with downregulation of contractile cytoskeleton proteins in the sarcomere and subsequent upregulation of cytoskeletal structural proteins, likely as a compensatory mechanism to bolster the cytoskeletal architecture. While we are unable to completely rule out changes in cardiomyocyte calcium handling as causative, these changes likely occur secondary to microtubule derangement and subsequent Jph2 redistributeion^8^. Rather than Sorbs2 acting as an RNA-binding protein in cardiomyocytes to regulate ion channels and ICD proteins, our findings here are consistent with the more established role for SORBS proteins as cytoskeletal crosslinking adapter proteins.

### 5.9 The SORBS Family and Clinical Genetic Associations

Our results indicate that Sorbs2 plays an essential role in maintaining normal cardiac function and is consistently dysregulated in clinical cardiomyopathy and rodent models of cardiac stress. To further explore the potential clinical relevance of Sorbs2, and other SORBS family members, we performed database and literature searches to assess if genetic variations in SORBS genes are associated with cardiac-related clinical phenotypes (results summarized in **Table 1)**. GWAS catalog search yielded a Sorbs2 variant associated with serum cardiac troponin T levels,^23^ a predictor of cardiovascular disease risk. Interestingly, this same variant is significantly associated with paroxysmal ventricular tachycardia in UK Biobank (UK-BB) GWAS data, summarized by PheWeb.^24^ With relevance to our findings in Sorbs2-cKO mice, query of the Cardiovascular Disease Knowledge Portal (CVD-KP) indicated strong associations for both Sorbs1 and Sorbs2 variants with P-wave interval/duration, the former of which was previously reported,^26^ and is also linked to cardiac arrest in UK-BB GWAS data. In addition, CVD-KP revealed a Sorbs2 variant associated with P-wave terminal force,^21^ a known indicator of LA enlargement, and this variant is significantly associated with DCM in published GWAS data^22^ and with decreased Sorbs2 expression in human cardiovascular tissue samples.^3^ While many of these associations are of sub-threshold significance by GWAS standards, in sum, these and other relevant associations highlighted in **Table 1** point to the interesting possibility that genetic variations in SORBS genes influence the onset and progression of several cardiovascular-related clinical phenotypes, with several notable instances that are consistent with our observations in Sorbs2-cKO mice (e.g., P-wave alterations, LA enlargement, and DCM).

## 6. Discussion

Beyond the field’s focus on the cytoskeletal/sarcomeric proteins themselves, this work highlights and reiterates an important role for cytoskeletal adapter proteins in the onset and development of cardiomyopathy. There has been mounting interest in the functions of SORBS proteins in cardiac biology and disease, yet their roles specifically in cardiomyocytes, where they are most highly expressed, has not been assessed using conditional gene deletion mouse models, or relevant transgene sequences (i.e., cardiac Sorbs2 transcript isoforms). To address this, we generated and characterized mice with cardiac-specific loss of Sorbs2, the most abundantly expressed SORBS family member in heart. In addition, we interrogated available bioinformatic datasets to examine Sorbs2 dysregulation in mouse models and patients with HF and whether SORBS genetic variants are associated with cardiac phenotypes. Overall, our studies provide key insights into the critical role for Sorbs2 in maintaining cardiac structure/function and highlight its potential clinical relevance.

In summary, Sorbs2 is consistently upregulated in humans with ischemic and idiopathic cardiomyopathies, and in experimental animal models of these diseases. Sorbs2 predominantly localizes to the intercalated disc and along sarcomeres at Z-discs, particularly adjacent to the lateral membrane at costameres in cardiomyocytes **(Figure 7)**. Sorbs2-cKO mice exhibit atrial and ventricular conduction defects, underlying diastolic dysfunction, develop progressive systolic dysfunction starting after 6 months of age, and die with congestive HF after 12 months of age. Systolic dysfunction is coincident with severely impaired cardiac contractility due in part to a failure to generate adequate mechanical tension in myofibers. Interrogation of cytoskeletal structures indicates that loss of Sorbs2 in cardiomyocytes does not significantly impair expression or distribution of ICD proteins, but instead leads to defective microtubule polymerization/stability and compensatory upregulation of structural proteins (alpha- and beta-tubulin, desmin, and vinculin) **(Figure 7)**. Our data, in conjunction with prior literature, support that Sorbs2 is an adapter protein that functions to maintain the structural integrity of the cardiomyocyte cytoskeleton by strengthening interactions between microtubules and other structural proteins at crosslink sites.

### 6.1 Microtubule and Cytoskeletal Proteins Relevant to Cardiac Function

Our work here may further direct research efforts to better understand how SORBS proteins regulate microtubule dynamics in cardiomyocytes, and beyond. Sorbs2 can bind to Tubb and enhance microtubule polymerization,^8, 37^ and our data complement these findings in showing that Sorbs2-cKO hearts have reduced levels of polymerized microtubules despite substantially increased tubulin protein levels, prior to systolic dysfunction. Other studies of age-related cardiomyopathies in mice reported decreased microtubule polymerization and aberrant microtubule dynamics coincident with HF.^39, 40^ Prior work also found increased microtubule abundance in human and animal models of dilated HF, ischemic cardiomyopathy and cardiac hypertrophy,^41–44^ with particular emphasis on elevations in detyrosinated Tuba, which increases binding of microtubule-associated proteins on plus ends, contributes to increased cell stiffness,^45, 46^ resisting both compressive and stretching forces.^47, 48^ Future work will be needed to determine if the microtubule changes observed in Sorbs2-cKO hearts are a primary consequence of Sorbs2 loss or a proximal adaptation to early systolic dysfunction. Although we observed changes in microtubule abundance in Sorbs2-cKO hearts prior to systolic dysfunction, suggesting a causal link, others have proposed that rewiring of microtubule and desmin networks may be an early-disease change prior to end-stage HF requiring transplant.^42^ While, we cannot be certain whether the microtubule changes are secondary to cytoskeletal weakening due to loss of Sorbs2 crosslinking, temporary downregulation of sarcomere components, or whether Sorbs2 directly modifies the microtubules, our data in conjunction with published data on SORBS proteins support reasonable speculation that these may underlie the onset of HF in Sorbs2-cKO mice.

Prior studies also indicate that the non-sarcomeric cytoskeleton is necessary for normal cardiomyocyte contractility, tension sensing and signal transduction.^49^ Considering the established role for SORBS proteins as actin filament crosslinking adaptor proteins, one interesting speculation is that Sorbs2 is critical for maintaining connections among the structural cytoskeleton (microtubules, Des, Actn2, Vcl, etc.) and the contractile cytoskeleton (sarcomeres) at cardiomyocyte z-discs, costameres, and ICDs. Recently, an immunoprecipitation mass spectrometry based proteomics approach was used to confirm Sorbs2 protein interactions with Tubb2a, Des, Actn1/2, Vcl, and Myh7/9 in heart and hESC-derived cardiomyocytes,^37^ perhaps suggesting a causal relationship between loss of Sorbs2 and dysregulation of these proteins.

Although we know these important connections are critical for coordinating cardiomyocytes excitation, contraction and relaxation mechanics, the molecular mechanisms for how individual cardiomyocytes sense changes in mechanical load and respond with compensatory changes in cytoskeletal proteins needs further investigation. We speculate that cardiomyocyte loss of Sorbs2 weakens the crosslinking interactions of tubulin, desmin, actin, and actinin proteins at z-discs and intercalated discs and that, over time (beat-by-beat), this manifests into detectable maladaptive remodeling of microtubule and intermediate filament networks, likely to increase tensile strength and modify cardiomyocyte contraction/relaxation. If so, this weakening is likely to occur to a very subtle extent (i.e., difficult to measure) given the delayed manifestation of contractile dysfunction observed in Sorbs2-cKO mice at about 6 months of age (>100,000,000 heart beats).

Additional published work strongly supports our speculation that Sorbs2 crosslinks the cardiomyocyte cytoskeleton through diverse protein: protein interactions (mediated by its SoHo and SH3 domains and proline rich motifs) with actin, microtubules, and associated proteins, as well as enabling signal transduction pathways. Sorbs2 was originally identified as an Abl2 (also known as ARG) interacting protein through a two-hybrid approach.^50^ This interaction has been reconfirmed and extends to other members of both the ABL and SORBS families and dozens of other Abl interacting proteins and phosphorylation targets including Abi1,^51^ Cbl,^52^ and others.^53^ ABL family kinases are necessary for cardiac growth and development,^54^ localize to focal adhesion and adherens junctions,^55, 56^ and exhibit multifaceted roles in the regulation of cytoskeleton proteins.^57^ In addition to their kinase activities, Abl1 and Abl2 have non-kinase dependent functional interactions with both actin fibers and microtubules that are sufficient to regulate the dynamics and stability of these cytoskeletal filaments.^56, 58, 59^ Future work is needed to determine how Sorbs2, which itself is phosphorylated by Abl kinases^50^ (as well as others), fits into Abl signal transduction pathways in cardiomyocytes. Beyond the Abl kinase family, other Sorbs2 interacting proteins have been identified with well described regulatory roles in cytoskeletal biology.^33^ The ubiquitin ligase Cbl is anchored by Sorbs2^60^ and directly regulates microtubule polymerization.^61^ Furthermore, large-scale protein-protein interaction studies suggests that Sorbs2 interacts with Mapre1 (also known as EB1),^62^ which belongs to a family of microtubule-end capping adaptor proteins that interact with SH3 domains through conserved proline-rich regions.^63^ Sorbs2 also interacts with the actin: myosin crosslinking protein Mybpc3;^64^ however, the relationship between these and other z-disc attachments and Sorbs2 interacting proteins^37^ remain unclear. Further and carefully designed experiments will be needed to define the cellular response to overexpression of Sorbs2 in mature cardiomyocytes as excess Sorbs2, above physiological levels, is sufficient to sequester interacting protein partners and collapse the cytoskeleton in cardiomyocytes^65^ as well as other cell types.^38^ This may confound the results observed when viral overexpression of Sorbs2 induced rapid HF in mice (i.e., within 3 weeks).^8^ Extensive future work will be needed to further tease apart the role for SORBS proteins in these dense cytoskeletal crosslinking networks, especially in mature cardiomyocytes.

### 6.2 Potential Mechanisms for Altered Cardiac Electrophysiology

Whole-body constitutive loss of Sorbs2 causes lethal ARVC in mice,^7^ and although some phenotypes are shared between the whole-body KO and cardiomyocyte-specific KO mice (i.e. bifid P-waves, QRS waveform anomalies, HF, death), notable discrepancies exist. Global Sorbs2-KO mice develop an aggressive “ARVC-like” phenotype, RV dilation, arrhythmias and die between ∼4-6 months of age. By contrast, Sorbs2-cKO mice show slow progressing DCM phenotype, without arrhythmias despite early LA enlargement, and die between ∼11-15 months-old. Sorbs2-cKO mice also do not show severe RV dilation nor clear signs of arrhythmia or altered Cx43 expression, which was astonishingly reduced by 90% in global Sorbs2 KO mice. Overall, global and cardiomyocyte-specific Sorbs2 KO mice are quite distinct, and our work helps to clarify the specific contribution of Sorbs2 loss in cardiomyocytes to cardiac structure and function in mice and highlights important likely roles for Sorbs2 in other cell types in contributing to diverse cardiac phenotypes.

Sorbs2-cKO mice exhibit p-wave alterations that may be due to increased atrial size rather than atrial conduction deficiency. Clinical presentation with bifid p-waves coincides with increased left atrial size,^31^ consistent with our observed correlations in Sorbs2-cKO. Given the profound cytoskeletal remodeling observed in Sorbs2-cKO mice, lack of disrupted Cx43 and NaV1.5 expression, and absence of atrial fibrillation, we speculate that the p-wave changes are predominantly of structural origin. Many potential physiological mechanisms could explain increased left atrial size including cardiac development, structural remodeling affecting atrial compliance (hypertrophy/fibrosis), or hemodynamic (left atrial pressure overload due to poor ventricular contractility). Future work will need to directly investigate the contribution of Sorbs2 expression in these and other atrial mechanisms.

Atrial-to-ventricular conduction defects (i.e., A-V block), or irregular heart rates were not present in Sorbs2-cKO mice but were in global KO mice. While the exact mechanism by which Sorbs2 controls cardiac conduction remains unknown; our data contest that this does not involve potential post-transcriptional regulation of Scn5a/NaV1.5, Kcnh2/mERG, or Gja1/Cx43 through Sorbs2 RNA-binding functions in cardiomyocytes, as suggested by prior global Sorbs2 KO mouse studies.^7, 29^ It is possible that the severe reduction of myocardial Gja1/Cx43 (and other ion channel expression and function) in global Sorbs2 KO mice occurs secondary to more severe HF, and not mechanistically related to Sorbs2 expression.^66^ This notion is further supported by the fact that Sorbs2 RNA-binding properties occur through its conserved ZnF-C2H2 domain located in human exon 35 (mouse exon 23);^67, 68^ and notably, this exon is restricted to alternate transcripts that are expressed in specific cell types (i.e. neuronal tissues) and largely absent from cardiomyocytes and heart tissues **(Supp. Figure 2a-b)**. It is worth reiterating that previous gain-of-function studies^29^ that demonstrate RNA-binding activity of Sorbs2 in cardiomyocytes utilized a neuronal Sorbs2 transcript harboring the RNA-binding domain.^9^ Beyond this, we further speculate that the more aggressive arrhythmia phenotypes in global Sorbs2 KO mice may result from contributions of key Sorbs2 functions in other cells and tissues (e.g., smooth muscle cells and neural regulation of cardiac functions), highlighting a subject of future study.

### 6.3 Clinical Relevance of Sorbs2 in Cardiac Disease

Precedent exists for various congenital cardiomyopathies to associate with Sorbs2 loss of function including atrial septal defects and transposition of arteries,^69, 70^ that may be due to c-Abl/Notch/Shh signaling in cardiac stem cells.^71^ Indeed, siRNA knockdown of Sorbs2 in cardiomyocyte differentiation studies decreases the expression of genes associated with cardiomyocyte maturation.^72^ While our work focuses on the slow progressing and adult onset of DCM, which likely results from subtle cytoskeletal changes/weakening, the possibility remains that subtle developmental alterations in Sorbs2-cKO mice could lead to slow progressing DCM.

Strong evidence supports that Sorbs2 is also associated with adult-onset heart disease, however the clinical significance of this remains unknown. Our interrogation of independent RNA expression data and western blot data shows consistent upregulation in Sorbs2 across a range of myocardial diseases, as has been recently noted.^73^ Further, Sorbs2 protein is upregulated in LVNC,^8^ diabetic cardiomyopathy,^6^ and is released from infarcted myocardium.^4^ Upregulation is potentially mediated by Mef2 transcription factors,^74^ although posttranscriptional regulation by disease-relevant microRNAs and RNA-binding proteins may also contribute.^5, 75, 76^ For example, Sorbs2 mRNA harbors strong interaction sites for miR-29 and miR-30, which both show decreased abundances in failing hearts.^77^ While it remains unknown if and how these different components may contribute to overall Sorbs2 isoform expressions in heart, we speculate that Sorbs2 is up-regulated in response to declining heart function, as cardiomyocytes attempt to increase the strength and stability of their cytoskeleton.

Beyond expression changes, our data queries also revealed several notable and relevant links between SORBS genetic variants and human phenotypes, including alterations in cardiac conduction and structure. While some of these associations do not exceed genome-wide significance, cumulatively, these links hint at the potential translational relevance of our mouse studies. Ultimately, more rigorous and targeted examinations in additional cohorts will be needed to clarify the potential clinical significance of these associations.

### 6.4 Additional Future Directions

Looking beyond Sorbs2, we have not resolved if other SORBS proteins initially compensate, at least partially and/or temporarily, for the loss of Sorbs2 in cardiomyocytes to maintain cardiac function. Future work will need to expand investigations into the involvement Sorbs1 and Sorbs3 in cardiac biology. Along with Sorbs2, both are expressed in myocyte and nonmyocyte populations in the heart, show dysregulated expression in disease, and contain genetic variants associated with various cardiovascular phenotypes. Beyond this, protein structures are highly conserved among the three family members, and all localize along sarcomeres, at costameres, and at intercalated discs in heart tissues. SORBS family proteins can interact with one another, share several other protein interaction partners, and provide some redundancy in biological systems. However, each protein can also display independent properties related to mechano-transduction.^78^ Considering that Sorbs1, Sorbs2, and Sorbs3 could collectively output at least 21 different protein isoforms, including several that are dysregulated in idiopathic or ischemic cardiomyopathy, it is clear that the known complexities of SORBS interactions and their contributions to cytoskeletal organization in cardiac biology and disease are only beginning to emerge.

## 7. Acknowledgements

Dr. Kenneth Margulies provided tissues through the Human Heart Tissue Bank at the University of Pennsylvania. Dr. Chad Grueter provided αMHC-Cre transgenic mice for use. Dr. Robert Weiss, Kathy Zimmerman, and Alyssa Bosko contributed to mouse echocardiographic analyses. Dr. Patrick Breheny advised on statistical analyses. Meg SmolikHagen contributed to mouse EKG analyses. Nathan Witmer and Gabrielle Abouassaly assisted with cloning and animal colony maintenance.

## Non-standard Abbreviations and Acronyms

αMHC: Cre alpha Myosin Heavy Chain promoter driving Cre-Recombinase
ARVC: Arrhythmogenic right ventricular cardiomyopathy
cKO: Cardiomyocyte-specific knockout
DOB: Dobutamine
dP/dT: Change in LV pressure over time
EF: Ejection Fraction
EGASXXX: European Genome-Phenome Archive Series Record
GSEXXX: Gene Expression Omnibus Series Record
HCM: Hypertrophic cardiomyopathy
HF: Heart Failure
ICD: Intercalated disc
LA: Left Atrium
LV: Left Ventricle
LVNC: Left ventricular non-compaction
NF: Non-failing hearts
NRCM: neonatal rat cardiomyocyte
pCa: Calcium concentration
PE: Phenylephrine
RV: Right Ventricle
RA: Right Atrium
TAC: Transverse Aortic Constriction
TPM: Transcripts Per Million
vg/g: viral vector genomes per gram of body weight
WGA: Wheat Germ Agglutinin

## SUPPLEMENTAL MATERIAL

### Supplemental Figure Legends

**Supplement Figure 1.**
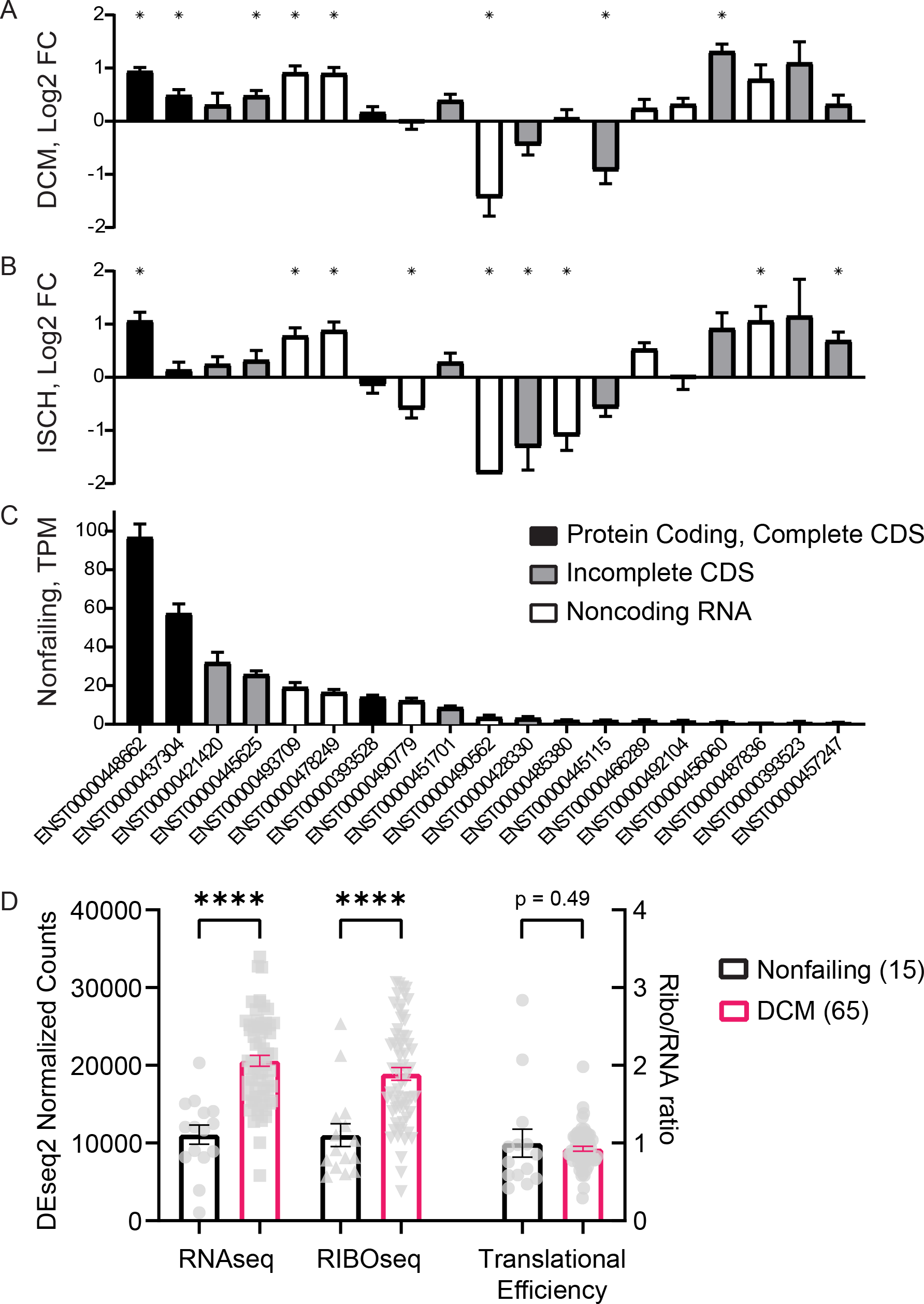
Transcriptional dysregulation of Sorbs2 in heart failure. (A-C) Human Sorbs2 transcript expression analysis of RNA sequencing data acquired from SRA archive PRJNA477855 remapped using the Kallisto/Sleuth pipeline. Sorbs2 transcript numbers are listed on X-axis in panel C. Black bars are high-confidence protein-coding transcripts with complete CDS, grey bars are transcripts with incomplete CDS, white bars are non-coding transcripts. Sample sizes are N=14 (Nonfailing), N=13 (Ischemic Cardiomyopathy, ISCH), N=37 (Dilated Cardiomyopathy, DCM), * p<0.05, compared to nonfailing. (A) Differential expression (Log2-fold-change) of transcripts in dilated cardiomyopathy. (B) Differential expression of Sorbs2 transcripts in Ischemic cardiomyopathy. (C) Average expression of Sorbs2 transcripts in non-failing hearts, sorted from most to least abundant with a lower cutoff of TPM>1, (represents 18 of 65 transcripts). (D) Left axis, RNA and ribosome reads, and on right axis translational efficiency (TE) plotted from public ribosomal profiling data in nonfailing and DCM human hearts from Van Heesch, et. al.^17^ Both RNA and Ribosome reads are significantly upregulated in DCM hearts, suggesting that cardiac Sorbs2 expression at baseline and disease is primarily under transcriptional rather than translational regulation (i.e., TE = 1 and is not significantly different in DCM).

**Supplement Figure 2.**
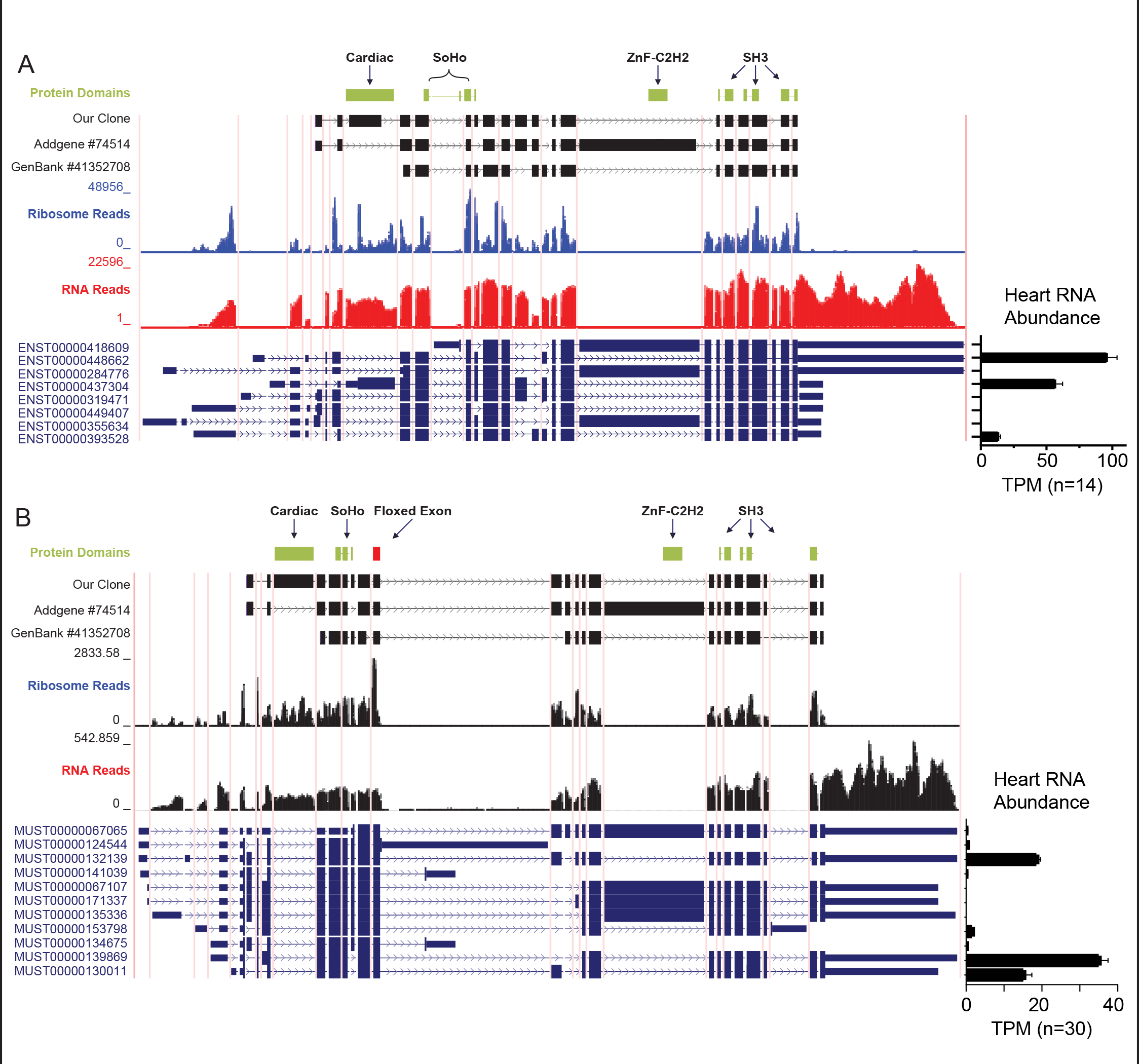
Sorbs2 genetic loci in human and mice (A) Sorbs2 genetic locus in humans. Vertical pink lines denote exon junctions. Green boxes denote the location of annotated protein domains, as well as the location of a cardiac-specific exon. Black boxes denote the alignment of three different SORBS2 clones published related to cardiac research. Blue (Ribosome) and Red (RNA) tracks show sequencing data from human hearts (adapted from Van Heesch, et.al ^17^). Protein coding transcript tracks list those with complete CDS annotations (Gencode Basic, 8 of 65 putative transcripts). To the right is transcript abundance in nonfailing human hearts based on RNA sequencing data described in **Supp. Figure 1C**. Abbreviations SoHo = Sorbin Homology Domain, SH3 = SRC Homology 3 Domain, ZnF-C2H2 = C2H2-type Zinc Finger (RNA-binding domain). Note this RNA-binding domain is encoded by Addgene plasmid #74514, which was used to show Sorbs2-RNA binding activity of cardiac ion channels ^29^, however it is located in an exon that is not expressed in human heart tissue, or present in two clones obtained from heart cDNA, including “our clone”, which was obtained by RT-PCR amplification of the cardiac-specific Sorbs2 isoform from mouse heart cDNA. (B) Sorbs2 genetic locus in mice, with a similar track layout. Note the red box indicating the obligate exon 12 with flanking loxP sites. These mice were crossed to αMHC- CRE expressing line to generate cardiomyocyte-specific Sorbs2 knockout mice (Sorbs2-cKO). The two tracks of black peaks represent Ribosome and RNA sequencing data specifically from cardiomyocytes, remapped from SRA-PRJNA484227 ^19^. Bar graph on right shows TPM expression of Sorbs2 protein coding transcripts (Gencode Basic, 11 of 37 putative transcripts) in mouse hearts from RNAseq data. ZnF-C2H2 represents the location of the identified RNA-binding domain in transcripts that are not abundant in hearts or cardiomyocytes. The cardiac-specific Sorbs2 exon remains unannotated in mouse reference transcriptomes yet is clearly present in RNA and Ribosome sequencing data.

**Supplement Figure 3.**
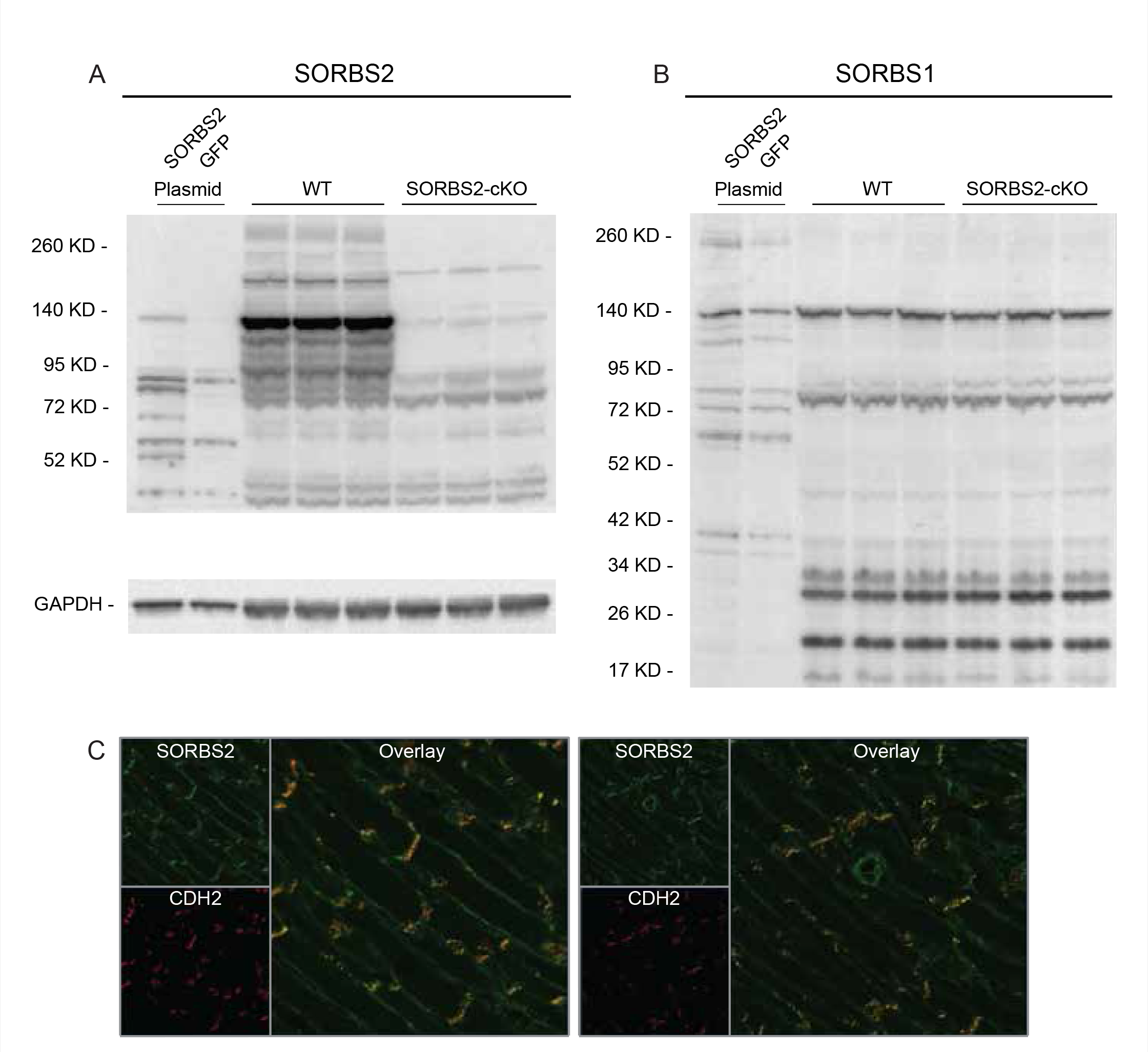
Cardiomyocyte-specific Sorbs2 knockout mice. (A) Western blot showing expression of Sorbs2, in WT and cKO heart samples from female mice at ∼12 months age using an independent antibody (Sigma mouse monoclonal). Also shown on left side is a positive control (Sorbs2 plasmid cloned from mouse heart cDNA expressed in HT1080 cells), as a confirmation of on-target antibody binding. (B) Western blot of Sorbs1 expression from WT and cKO heart tissues does not show compensatory changes in Sorbs1 expression following cardiomyocyte loss of Sorbs2. Gapdh shown as loading control for panels A and B. (C) Representative Sorbs2 immunofluorescence (green) in heart sections from wildtype (WT) mice, co-localized with N-Cadherin (CDH2, red) at the intercalated disc (ICD). Also note Sorbs2 reactivity in the coronary artery on the right picture.

**Supplement Figure 4.**
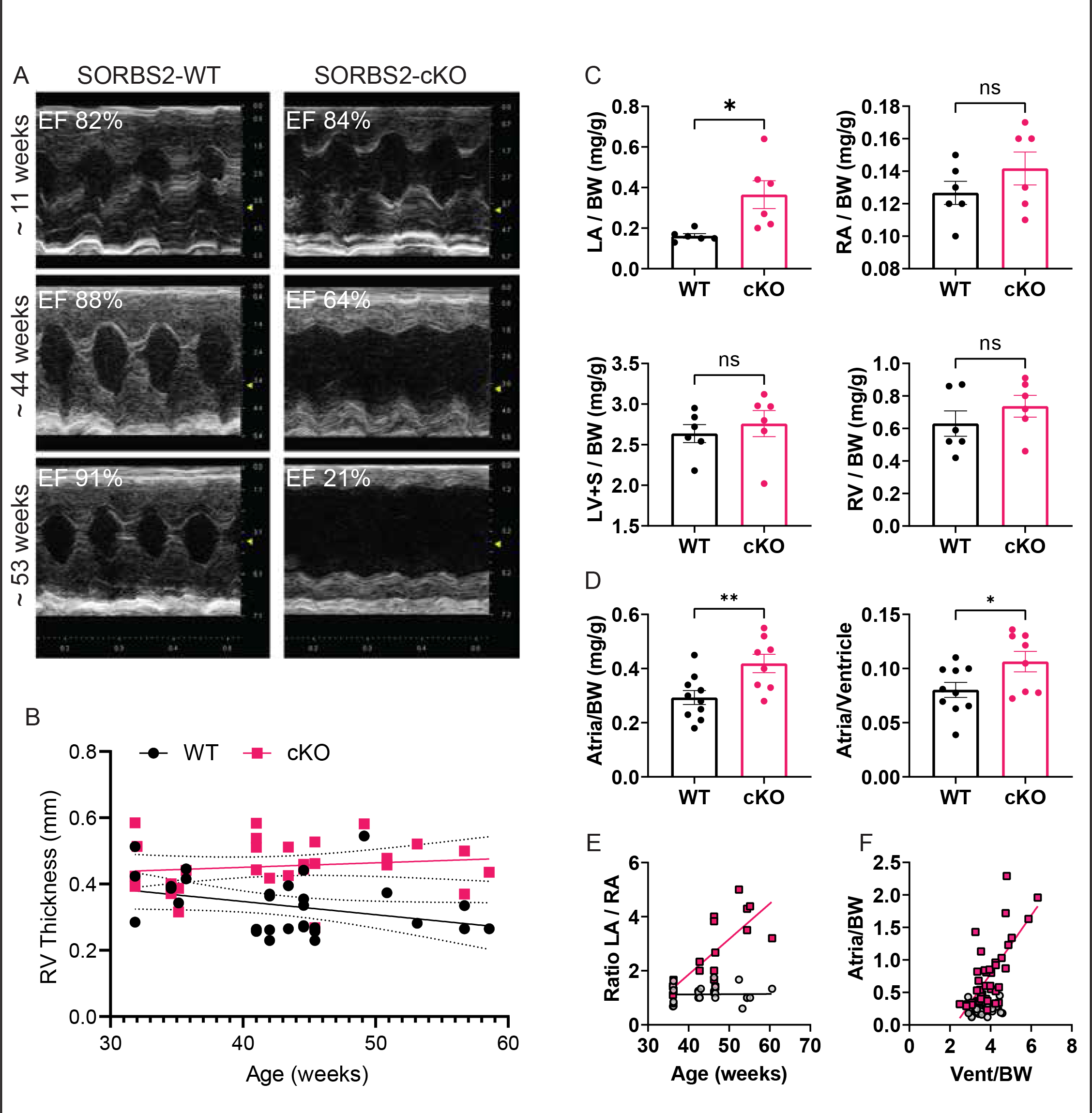
Cardiomyocyte-specific Sorbs2 knockout mice develop age-related systolic dysfunction, cardiac remodeling, and premature death. (A) Representative serial m-mode echocardiographic images from one WT and one Sorbs2-cKO mice. The callout indicates the ejection fraction (EF) for that mouse calculated using the Endocardial and Epicardial traces (see Methods section 2.3). At three months old (first row) both WT and cKO mice show normal wall motion, wall thickness, and chamber dimensions in systole and diastole. At 10 months (second row) Sorbs2-cKO heart shows reduced wall motion, increased chamber dimension, and decreased EF. At 12 months old, (third row) Sorbs2-cKO hearts show significant LV systolic dysfunction with a big chamber, thin walls, and poor wall motion. (B) Time course of echocardiography from WT and Sorbs2-cKO hearts indicates trending to increased RV thickness in cKO hearts, coincident with emerging HF. (C) Postmortem gravimetric analyses of heart chamber mass in ∼48-week male mice, normalized to bodyweight (BW). The left atrium (LA) is significantly enlarged in Sorbs2-cKO hearts compared to control, whereas, the right atrium (RA), the left ventricle plus septum (LV+S) and the right ventricle (RV) are not different. Dots show individual hearts, N=6 per group, with mean +/- SEM. Significance from t-test, * p<0.05, ns = not significant. (D) Postmortem gravimetric analysis of atrial mass normalized to either body weight (BW) or ventricular mass in mice 12-24 weeks of age Dots show individual hearts with mean +/- SEM. Significance from t-test, * p<0.05, **p<0.01. (E) Male ratio of postmortem LA/RA mass over time (cKO regression, r^2^=0.596, p=2e^-4^). This indicates increases in LA mass rather than RA or bi-atrial mass are driving the significant increase in total atrial mass shown in Figure 3D. (F) Male correlation of atrial to ventricle size in WT and cKO mice hearts (cKO regression, r^2^=0.479, p<1e^-4^).

**Supplement Figure 5.**
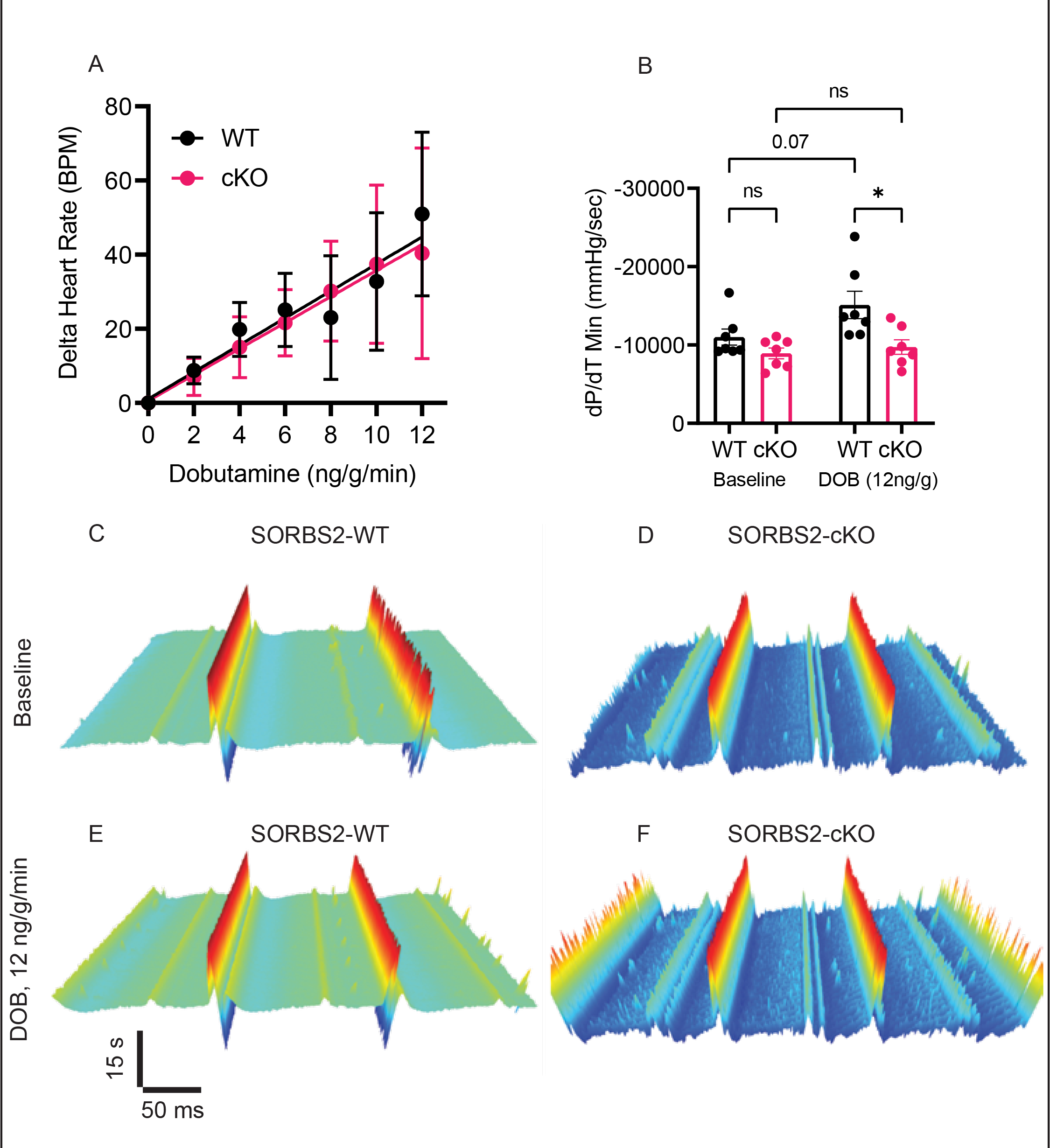
Cardiomyocyte-specific Sorbs2 knockout mice have severe contractile dysfunction. (A) Datapoints represent the group mean +/- SEM change in heart rate from baseline through increasing dobutamine (DOB) concentrations (N=7 mice per group), and a black trendline denotes a linear regression fit from baseline through the highest dose (12 ng/g/min). No difference was found. These data show that Sorbs2-cKO hearts are unable to increase cardiac contractility in response to DOB despite having equivalent increases in heart rate. (B) Final 30 second average per dose for dP/dT Min. Dots show individual mice with mean +/- SEM, statistics acquired using one-way ANOVA with Sidak posthoc test comparing selected groups (each comparison shown on plot), * p<0.05, ns = not significant. (C-F) Waterfall plots of EKG intervals (30 seconds stacked back to front) in WT and Sorbs2-cKO mice. (C) WT hearts have normal sinus rhythm with regular R-R, P-P, and P-R intervals. (D) cKO hearts also show normal sinus rhythm and intervals, despite obvious bifid P-waves, increased P duration, and QRS duration. (E-F) WT and cKO hearts after DOB challenge, both have expected increase in heart rate, indicated by reduced R-R interval, and maintain sinus rhythm.

**Supplement Figure 6.**
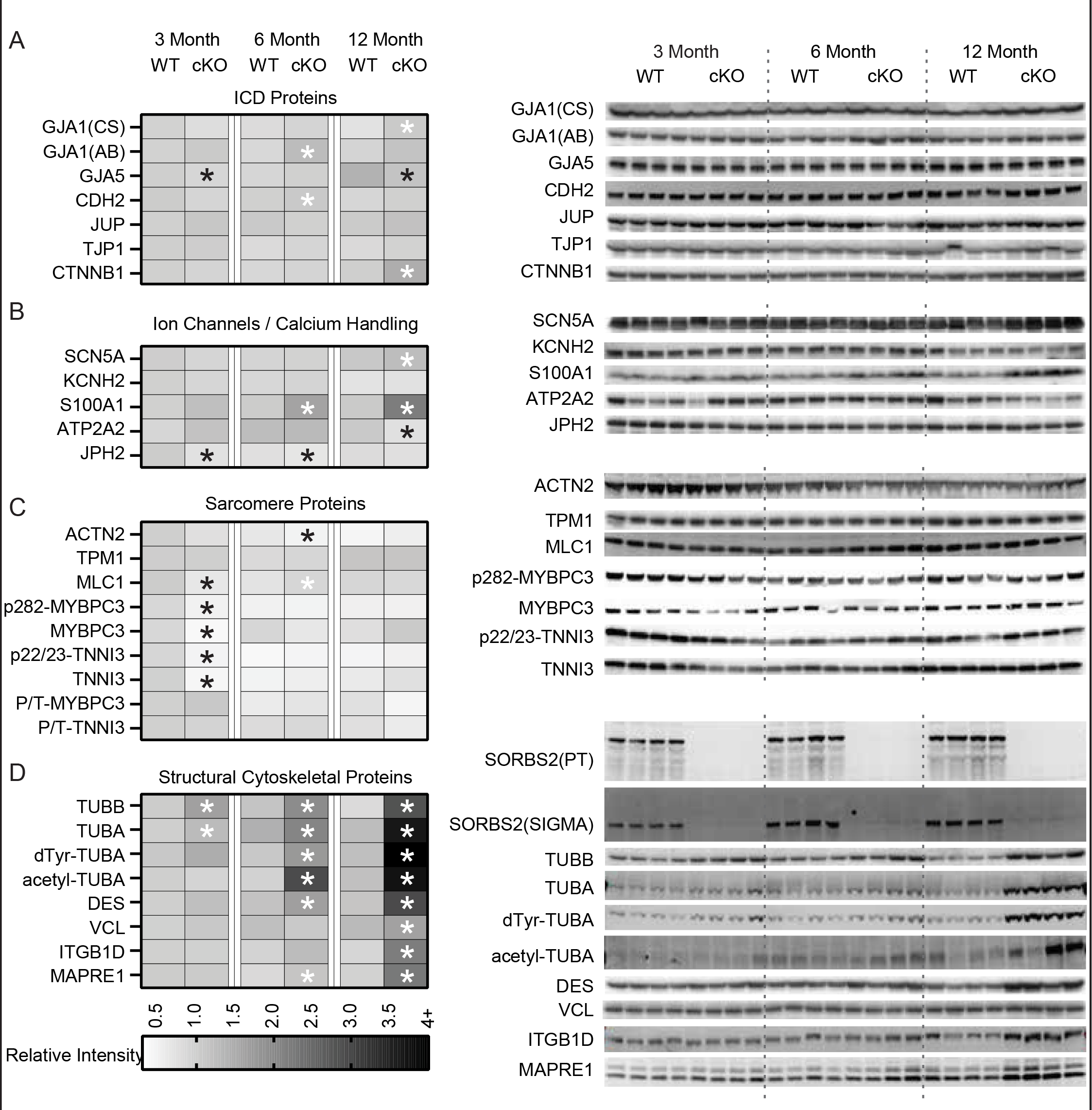
Cardiomyocyte-specific Sorbs2 knockout mice have dysregulated cytoskeletal protein expression not ICD. (A-D) Heatmaps reprint from Figure 6 alongside digitized western blot images that show analysis of protein expression from WT and Sorbs2-cKO cardiac lysates at 3, 6, and 12 months of age. Each box represents the mean integrated intensity, normalized to loading control and expressed relative to 3-month WT; n=4 mice per group. Significant differences (p<0.05) denoted with (*, black for down-regulated and white for up-regulated) overlay on the heatmap, were acquired using t-test comparing cKO to WT at each age. (Abbreviations: CS = Cell Signaling, AB = Abclonal). Heatmaps and blots are organized by (A) intercalated disc (ICD) proteins, (B) ion channels and calcium handling proteins, (C) sarcomere proteins, and (D) structural cytoskeletal proteins.

**Supplement Figure 7.**
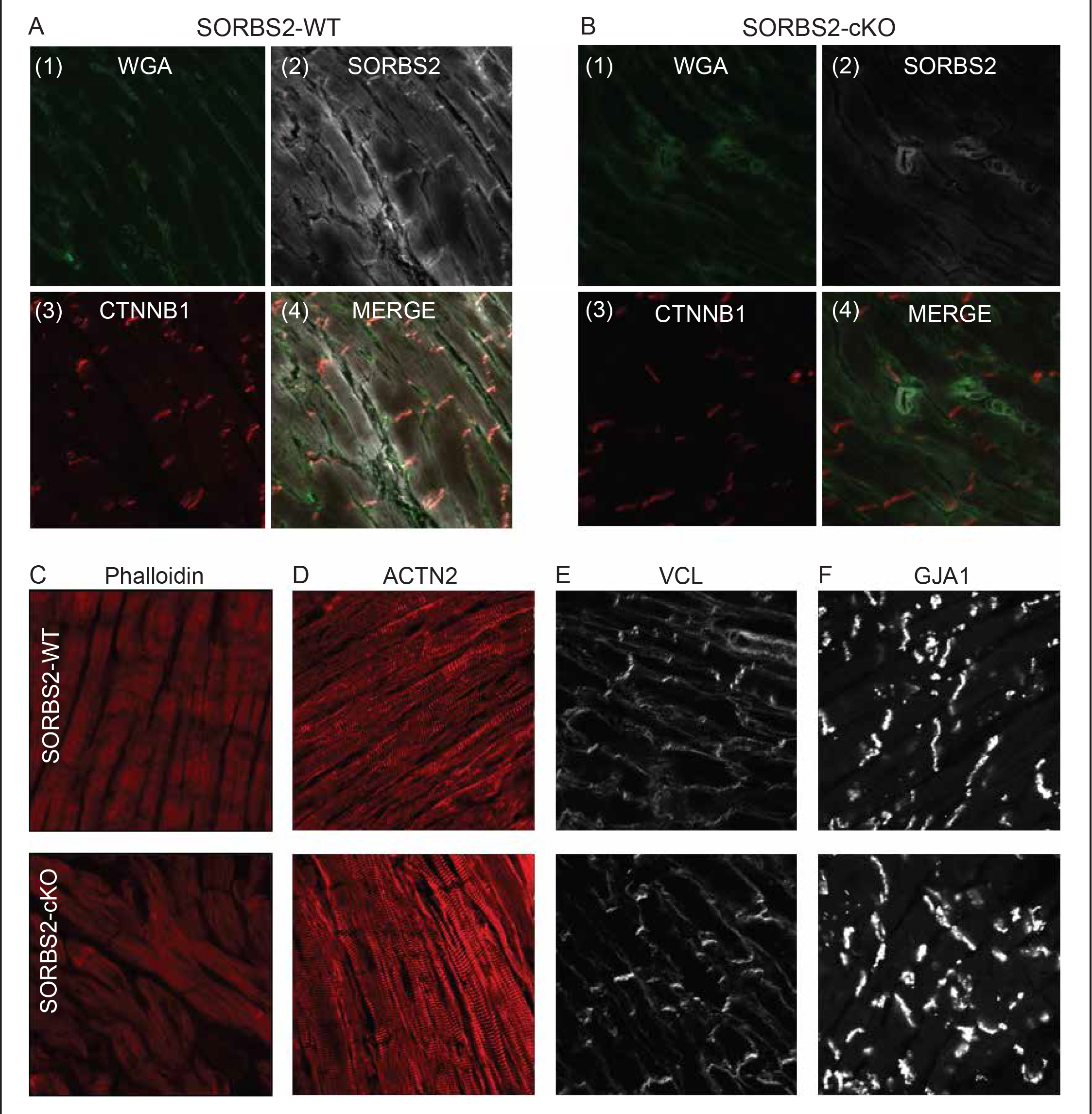
Cardiomyocyte-specific Sorbs2 knockout mice have normal localization of cytoskeletal and ICD proteins. (A-B) Representative immunofluorescent microscope images of heart tissue oriented with longitudinal myofibers from (A) WT and (B) Sorbs2-cKO mice. Inset pictures show staining with (1) wheat germ agglutinin (WGA) in green, (2) Sorbs2 in white, (3) Ctnnb1 in red, and (4) a color overlay. A blinded review of images from WT and cKO mice for Ctnnb1 localization and several other cytoskeletal and ICD proteins (C-F) were indistinguishable from each other, (n=4 images per mouse and n=4 mice per group). These include (C) phalloidin staining actin fibers in red, (D) alpha actinin 2 (Actn2) staining Z-disk and sarcomeres in red, (E) vinculin (Vcl) staining ICDs and costameres in white, and (F) connexin 43 (Gja1) staining ICDs and vasculature in white. Representative WT images shown on the top row with cKO images on the bottom row.

**Supplement Figure 8.**
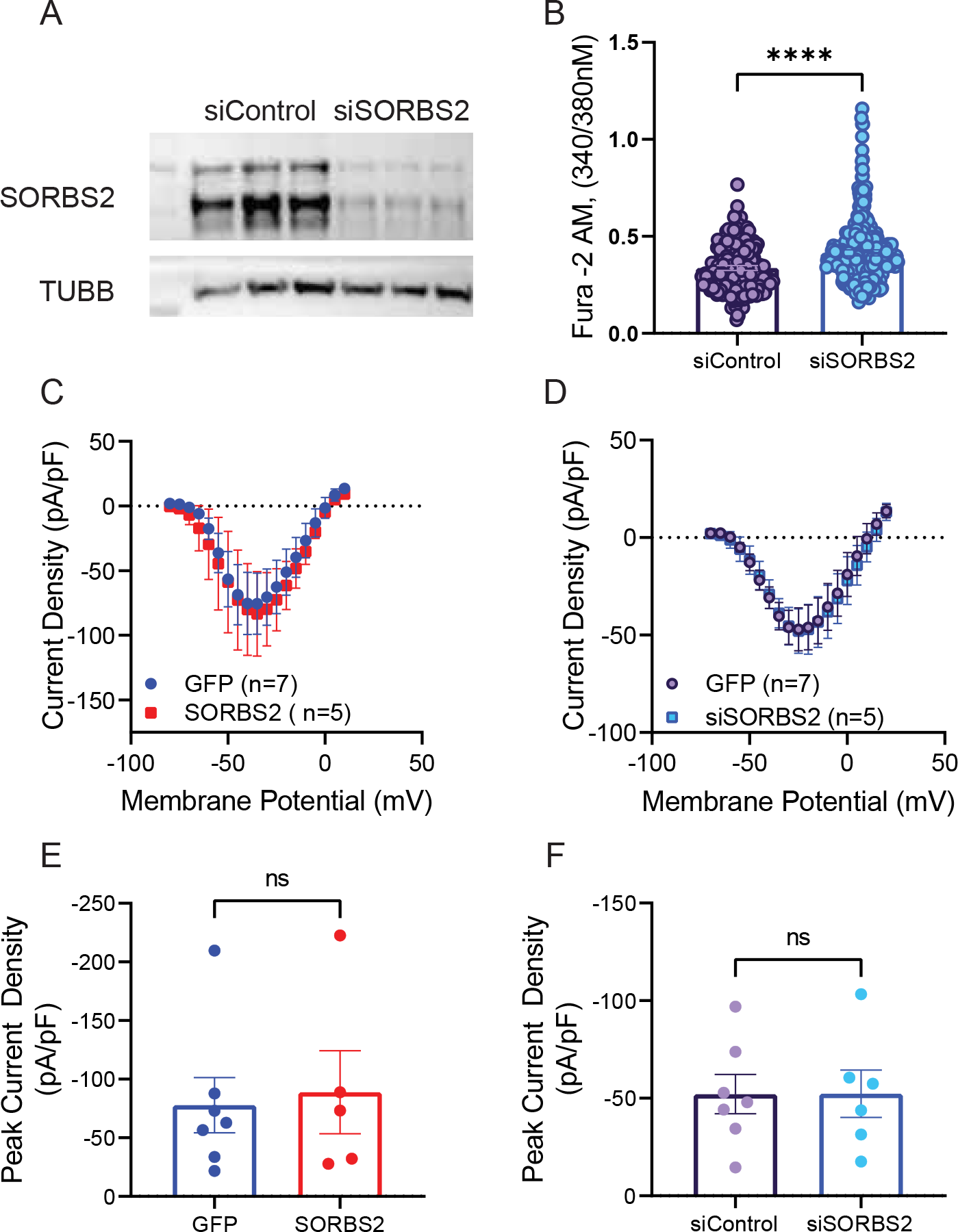
Sorbs2 regulation of intracellular calcium and Scn5a activity *in vitro*. (A) Western blot analysis of Sorbs2 protein levels in NRCMs transiently transfected with control or Sorbs2 siRNAs for ∼48 hours. TUBB shows consistent loading. (B) Intracellular calcium signal intensity from Fura-2AM loaded NRCMs. Data were acquired in 4 independent experiments, 2 plates per experiment per condition with ∼ 40-50 cells analyzed per plate. Dots represent individual cells with mean +/- SEM, statistics are acquired from T-test, **** p<0.0001, ns=not significant. (C-D) Overexpression or knock down of Sorbs2 did not significantly alter sodium current-voltage relationships derived from electrophysiology experiments in transiently transfected NRCMs. (E-F) Peak current density is also not significantly different. Data shown are mean SEM, with individual cells plotted in (D, E), N = 5-7 cells per treatment, statistics are acquired from T-test, ns = not significant.

### Supplemental Tables and Supporting Information

**Supplement Table 1.**
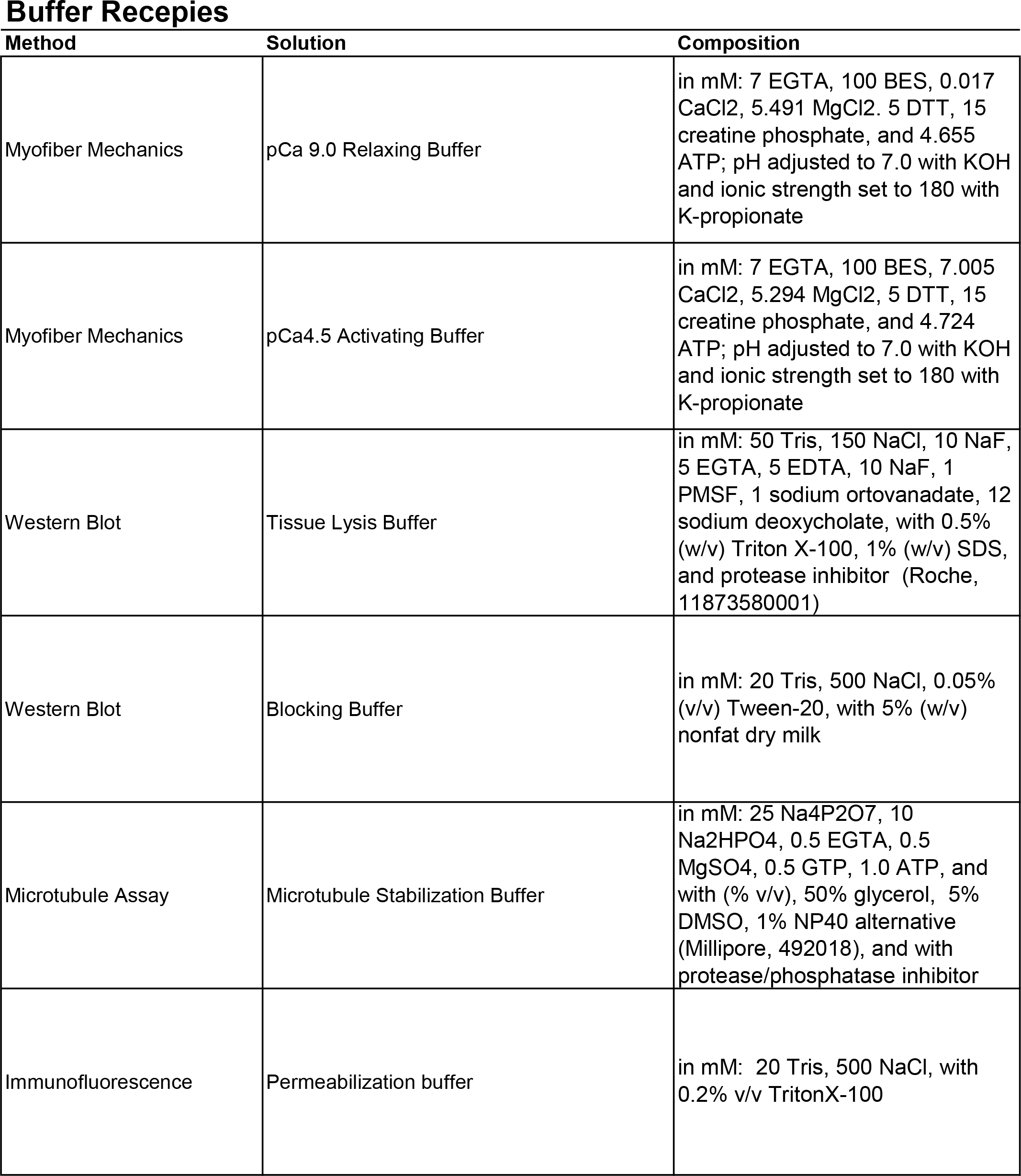
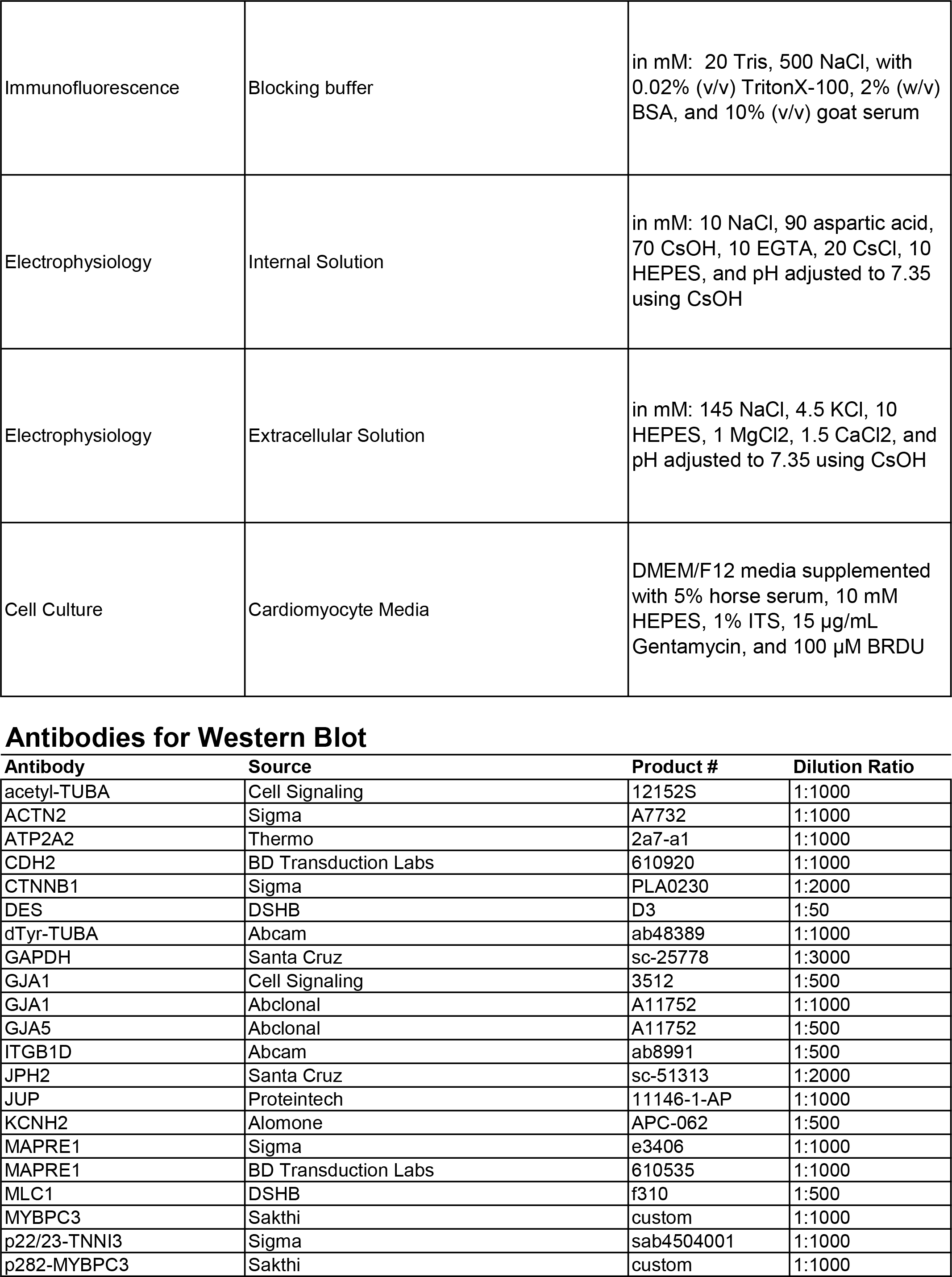
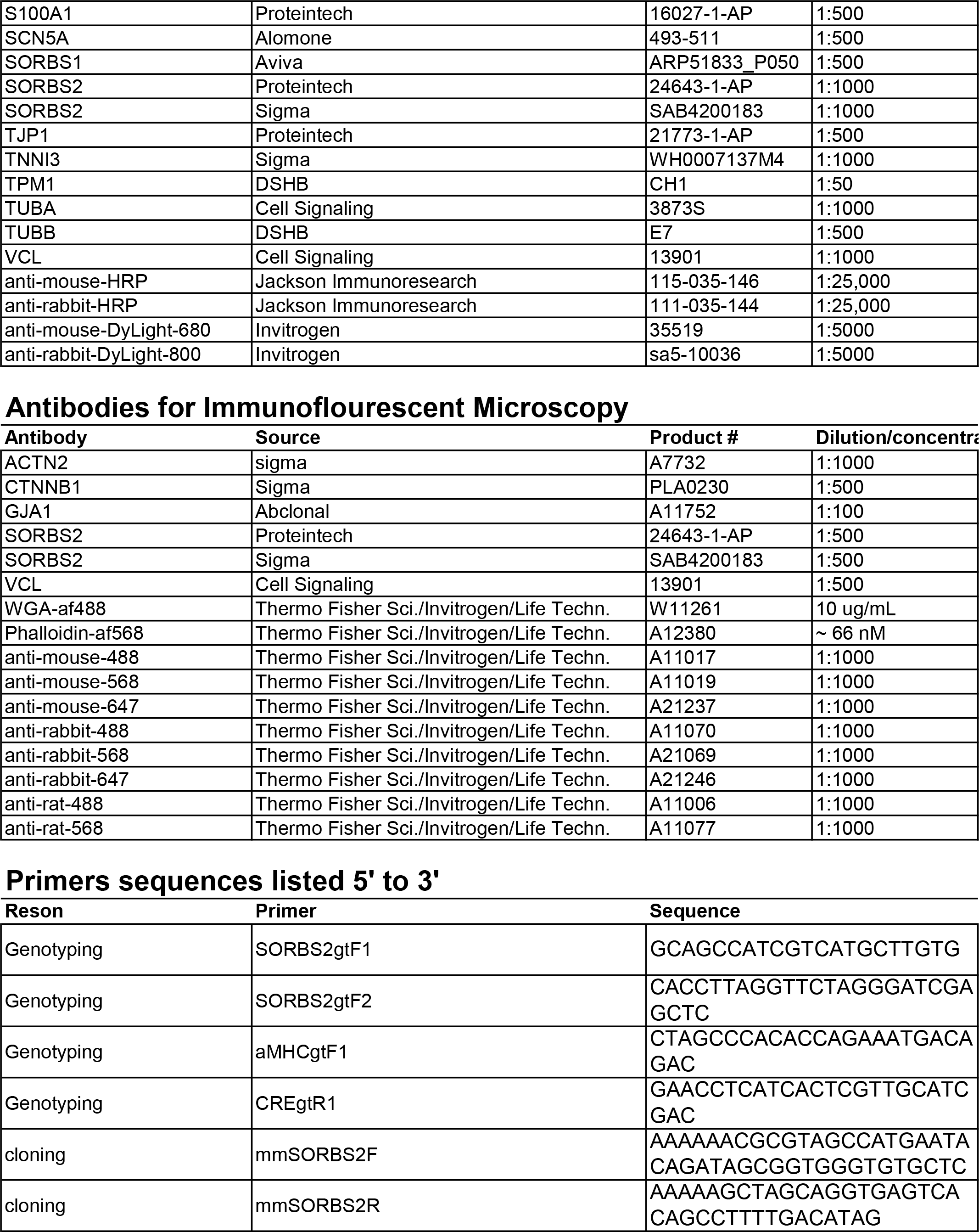
Supplement to Materials and Methods.

**Supplement Table 2.** Dataset description for Sorbs2 RNA dysregulation in heart failure. Table includes description for datasets used in Figure 1a. Information includes the accession number (Acc#), disease, sample number for control and disease groups, log2-fold-change value (l2fc) and -log10 p-value (-l10P). Blank rows correspond to the gaps in the Figure 1a plot.

| Acc# | Disease | Control N | Disease N | I2fc | -I10P |
| --- | --- | --- | --- | --- | --- |
| GSE36961 | HCM | 39 | 106 | 0.827 | 31.157 |
| GSE29819 | ARVC | 6 | 6 | 0.466 | 3.623 |
| GSE57345 | Idiopathic | 139 | 84 | 0.414 | 26.111 |
| EGAS00001002454 | DCM | 113 | 149 | 0.290 | 25.738 |
| GSE3586 | DCM | 15 | 13 | 1.009 | 7.438 |
| GSE5406 | Idiopathic | 16 | 86 | 0.677 | 5.300 |
| GSE1145 | Idiopathic | 11 | 15 | 1.454 | 5.126 |
| GSE79962 | HF | 11 | 9 | 0.416 | 3.876 |
| GSE46224 | HF | 8 | 8 | 0.650 | 2.988 |
| GSE53081 | Idiopathic | 4 | 8 | 0.362 | 1.166 |
| GSE57345 | Ischemic | 139 | 96 | 0.254 | 6.767 |
| GSE1145 | Ischemic | 11 | 11 | 1.595 | 3.699 |
| GSE5406 | Ischemic | 16 | 108 | 0.463 | 2.062 |
| GSE53081 | Ischemic | 4 | 8 | 0.533 | 2.023 |
| GSE16499 | Ischemic | 15 | 15 | 0.264 | 0.903 |
| GSE48166 | Ischemic | 15 | 15 | 0.372 | 0.210 |
| GSE13874 | genetic HF | 4 | 4 | 1.565 | 5.903 |
| GSE68518 | genetic HF | 6 | 6 | 0.554 | 4.559 |
| GSE36074 | TAC/HF | 5 | 7 | 0.596 | 5.712 |
| GSE58455 | TAC | 4 | 4 | 0.557 | 4.413 |
| GSE36074 | TAC/NF | 5 | 7 | 0.368 | 3.285 |
| GSE4648 | MI | 10 | 12 | 0.884 | 2.526 |
| GSE107569 | TAC | 6 | 6 | 0.786 | 2.151 |
| GSE35350 | TAC | 4 | 3 | 0.670 | 2.025 |
| GSE5500 | TAC | 4 | 6 | 0.384 | 1.290 |
| GSE5996 | cells Strech | 2 | 2 | 0.602 | 5.648 |
| GSE5996 | cells PE | 2 | 2 | 0.587 | 3.358 |
| GSE73896 | cells-PE | 4 | 4 | 0.524 | 2.770 |

Ejection Fraction
| Bin (weeks) | WT (mean) | cKO (mean) | (pval) |
| --- | --- | --- | --- |
| 10-20 | 0.793 | 0.765 | 0.362 |
| 20-30 | 0.883 | 0.847 | 0.067 |
| 30-40 | 0.798 | 0.687 | 0.035 |
| 40-50 | 0.832 | 0.596 | 9.48E-09 |
| 50-60 | 0.840 | 0.419 | 7.23E-21 |

Indexed EDV
| Bin (weeks) | WT (mean) | cKO (mean) | (pval) |
| --- | --- | --- | --- |
| 10-20 | 0.365 | 0.377 | 0.735 |
| 20-30 | 0.374 | 0.418 | 0.301 |
| 30-40 | 0.422 | 0.483 | 0.120 |
| 40-50 | 0.369 | 0.495 | 3.07E-04 |
| 50-60 | 0.389 | 0.734 | 9.54E-11 |

LV Mass
| Bin (weeks) | WT (mean) | cKO (mean) | (pval) |
| --- | --- | --- | --- |
| 10-20 | 60.067 | 59.129 | 0.826 |
| 20-30 | 71.044 | 69.945 | 0.795 |
| 30-40 | 95.392 | 96.634 | 0.763 |
| 40-50 | 82.019 | 85.332 | 0.421 |
| 50-60 | 78.739 | 95.106 | 0.001 |

LV Thickness
| Bin (weeks) | WT (mean) | cKO (mean) | (pval) |
| --- | --- | --- | --- |
| 10-20 | 0.756 | 0.760 | 0.911 |
| 20-30 | 0.772 | 0.746 | 0.638 |
| 30-40 | 0.841 | 0.794 | 0.131 |
| 40-50 | 0.832 | 0.862 | 0.238 |
| 50-60 | 0.841 | 0.777 | 0.018 |

Atria / BW
| Bin (weeks) | WT (mean) | cKO (mean) | (pval) |
| --- | --- | --- | --- |
| 10-20 | 0.270 | 0.397 | 0.094 |
| 20-30 | 0.328 | 0.427 | 0.138 |
| 30-40 | 0.218 | 0.335 | 1.03E-04 |
| 40-50 | 0.256 | 0.496 | 4.29E-05 |
| 50-60 | 0.208 | 0.982 | 1.87E-08 |
| 60-70 | 0.319 | 1.058 | 0.022 |

Ventricle / BW
| Bin (weeks) | WT (mean) | cKO (mean) | (pval) |
| --- | --- | --- | --- |
| 10-20 | 3.758 | 3.965 | 0.560 |
| 20-30 | 3.654 | 3.955 | 0.448 |
| 30-40 | 3.354 | 3.164 | 0.219 |
| 40-50 | 3.372 | 3.502 | 0.515 |
| 50-60 | 3.377 | 4.298 | 1.74E-05 |
| 60-70 | 3.252 | 3.623 | 0.377 |

**Supplement Table 3.** Summary statistics for cardiomyocyte-specific Sorbs2 knockout mice. Table supplements data plotted in Figure 4. Mean values were binned on weeks on age for WT and Sorbs2-cKO mice. P-values indicate difference between WT and cKO curves analyzed using 2-way ANOVA with Sidak’s multiple comparisons testing the interaction between age and genotype.

## References

1. Ntalla I, Weng LC, Cartwright JH, et al. Multi-ancestry gwas of the electrocardiographic pr interval identifies 202 loci underlying cardiac conduction. Nat Commun. 2020;11:2542

2. Carithers LJ, Ardlie K, Barcus M, et al. A novel approach to high-quality postmortem tissue procurement: The gtex project. Biopreserv Biobank. 2015;13:311–319

3. Aguet F, Brown AA, Castel SE, et al. Genetic effects on gene expression across human tissues. Nature. 2017;550:204–213

4. Kakimoto Y, Ito S, Abiru H, Kotani H, Ozeki M, Tamaki K, Tsuruyama T. Sorbin and sh3 domain- containing protein 2 is released from infarcted heart in the very early phase: Proteomic analysis of cardiac tissues from patients. J Am Heart Assoc. 2013;2:e000565

5. Bang C, Batkai S, Dangwal S, et al. Cardiac fibroblast-derived microrna passenger strand- enriched exosomes mediate cardiomyocyte hypertrophy. J Clin Invest. 2014;124:2136–2146

6. Shao M, Chen J, Zheng S. Comparative proteomics analysis of myocardium in mouse model of diabetic cardiomyopathy using the itraq technique. Adv Clin Exp Med. 2018;27:1469–1475

7. Ding Y, Yang J, Chen P, et al. Knockout of sorbs2 protein disrupts the structural integrity of intercalated disc and manifests features of arrhythmogenic cardiomyopathy. J Am Heart Assoc. 2020;9:e017055

8. Li C, Liu F, Liu S, et al. Elevated myocardial sorbs2 and the underlying implications in left ventricular noncompaction cardiomyopathy. EBioMedicine. 2020;53:102695

9. Zhang Q, Gao X, Li C, et al. Impaired dendritic development and memory in sorbs2 knock-out mice. J Neurosci. 2016;36:2247–2260

10. Agah R, Frenkel PA, French BA, Michael LH, Overbeek PA, Schneider MD. Gene recombination in postmitotic cells. Targeted expression of cre recombinase provokes cardiac-restricted, site- specific rearrangement in adult ventricular muscle in vivo. J Clin Invest. 1997;100:169–179

11. Spitler KM, Ponce JM, Oudit GY, Hall DD, Grueter CE. Cardiac med1 deletion promotes early lethality, cardiac remodeling, and transcriptional reprogramming. Am J Physiol Heart Circ Physiol. 2017;312:H768–h780

12. Hu P, Zhang D, Swenson L, Chakrabarti G, Abel ED, Litwin SE. Minimally invasive aortic banding in mice: Effects of altered cardiomyocyte insulin signaling during pressure overload. Am J Physiol Heart Circ Physiol. 2003;285:H1261–1269

13. Calligaris SD, Ricca M, Conget P. Cardiac stress test induced by dobutamine and monitored by cardiac catheterization in mice. J Vis Exp. 2013

14. Lynch TLt, Kumar M, McNamara JW, et al. Amino terminus of cardiac myosin binding protein-c regulates cardiac contractility. J Mol Cell Cardiol. 2021;156:33–44

15. Golden H, Gollapudi D, Gerilechaogetu F, Li J, Cristales R, Peng X, Dostal D. Isolation of cardiac myocytes and fibroblasts from neonatal rat pups. Methods in molecular biology (Clifton, N.J.). 2012;843:205–214

16. Matasic DS, Yoon JY, McLendon JM, et al. Modulation of the cardiac sodium channel na(v)1.5 peak and late currents by nad(+) precursors. J Mol Cell Cardiol. 2020;141:70–81

17. van Heesch S, Witte F, Schneider-Lunitz V, et al. The translational landscape of the human heart. Cell. 2019;178:242–260.e229

18. Sweet ME, Cocciolo A, Slavov D, et al. Transcriptome analysis of human heart failure reveals dysregulated cell adhesion in dilated cardiomyopathy and activated immune pathways in ischemic heart failure. BMC Genomics. 2018;19:812

19. Doroudgar S, Hofmann C, Boileau E, et al. Monitoring cell-type-specific gene expression using ribosome profiling in vivo during cardiac hemodynamic stress. Circulation Research. 2019;125:431–448

20. Buniello A, MacArthur JAL, Cerezo M, et al. The nhgri-ebi gwas catalog of published genome- wide association studies, targeted arrays and summary statistics 2019. Nucleic Acids Res. 2019;47:D1005–d1012

21. Christophersen IE, Magnani JW, Yin X, et al. Fifteen genetic loci associated with the electrocardiographic p wave. Circ Cardiovasc Genet. 2017;10

22. Esslinger U, Garnier S, Korniat A, et al. Exome-wide association study reveals novel susceptibility genes to sporadic dilated cardiomyopathy. PLoS One. 2017;12:e0172995

23. Welsh P, Preiss D, Hayward C, et al. Cardiac troponin t and troponin i in the general population. Circulation. 2019;139:2754–2764

24. Gagliano Taliun SA, VandeHaar P, Boughton AP, et al. Exploring and visualizing large-scale genetic associations by using pheweb. Nat Genet. 2020;52:550–552

25. Murray SS, Smith EN, Villarasa N, et al. Genome-wide association of implantable cardioverter- defibrillator activation with life-threatening arrhythmias. PLoS One. 2012;7:e25387

26. van Setten J, Brody JA, Jamshidi Y, et al. Pr interval genome-wide association meta-analysis identifies 50 loci associated with atrial and atrioventricular electrical activity. Nat Commun. 2018;9:2904

27. Ashar FN, Mitchell RN, Albert CM, et al. A comprehensive evaluation of the genetic architecture of sudden cardiac arrest. Eur Heart J. 2018;39:3961–3969

28. Kichaev G, Bhatia G, Loh PR, Gazal S, Burch K, Freund MK, Schoech A, Pasaniuc B, Price AL. Leveraging polygenic functional enrichment to improve gwas power. Am J Hum Genet. 2019;104:65–75

29. Qian LL, Sun X, Yang J, Wang XL, Ackerman MJ, Wang RX, Xu X, Lee HC, Lu T. Changes in ion channel expression and function associated with cardiac arrhythmogenic remodeling by sorbs2. Biochim Biophys Acta Mol Basis Dis. 2021;1867:166247

30. Pugach EK, Richmond PA, Azofeifa JG, Dowell RD, Leinwand LA. Prolonged cre expression driven by the α-myosin heavy chain promoter can be cardiotoxic. J Mol Cell Cardiol. 2015;86:54–61

31. Waggoner AD, Adyanthaya AV, Quinones MA, Alexander JK. Left atrial enlargement. Echocardiographic assessment of electrocardiographic criteria. Circulation. 1976;54:553–557

32. Ruan Y, Liu N, Priori SG. Sodium channel mutations and arrhythmias. Nat Rev Cardiol. 2009;6:337–348

33. Kioka N, Ueda K, Amachi T. Vinexin, cap/ponsin, argbp2: A novel adaptor protein family regulating cytoskeletal organization and signal transduction. Cell Struct Funct. 2002;27:1–7

34. Hallock PT, Chin S, Blais S, Neubert TA, Glass DJ. Sorbs1 and -2 interact with crkl and are required for acetylcholine receptor cluster formation. Mol Cell Biol. 2016;36:262–270

35. Pleger ST, Shan C, Ksienzyk J, et al. Cardiac aav9-s100a1 gene therapy rescues post-ischemic heart failure in a preclinical large animal model. Sci Transl Med. 2011;3:92ra64

36. Most P, Remppis A, Pleger ST, Katus HA, Koch WJ. S100a1: A novel inotropic regulator of cardiac performance. Transition from molecular physiology to pathophysiological relevance. Am J Physiol Regul Integr Comp Physiol. 2007;293:R568–577

37. Li C, Zheng Y, Liu Y, Jin GH, Pan H, Yin F, Wu J. The interaction protein of sorbs2 in myocardial tissue to find out the pathogenic mechanism of lvnc disease. Aging (Albany NY). 2022;14:800–810

38. Fredriksson-Lidman K, Van Itallie CM, Tietgens AJ, Anderson JM. Sorbin and sh3 domain- containing protein 2 (sorbs2) is a component of the acto-myosin ring at the apical junctional complex in epithelial cells. PLoS One. 2017;12:e0185448

39. Li L, Zhang Q, Zhang X, et al. Microtubule associated protein 4 phosphorylation leads to pathological cardiac remodeling in mice. EBioMedicine. 2018;37:221–235

40. Yashirogi S, Nagao T, Nishida Y, et al. Ampk regulates cell shape of cardiomyocytes by modulating turnover of microtubules through clip-170. EMBO Rep. 2021;22:e50949

41. Chen CY, Caporizzo MA, Bedi K, et al. Suppression of detyrosinated microtubules improves cardiomyocyte function in human heart failure. Nat Med. 2018;24:1225–1233

42. Schuldt M, Pei J, Harakalova M, et al. Proteomic and functional studies reveal detyrosinated tubulin as treatment target in sarcomere mutation-induced hypertrophic cardiomyopathy. Circ Heart Fail. 2021;14:e007022

43. Belmadani S, Poüs C, Ventura-Clapier R, Fischmeister R, Méry PF. Post-translational modifications of cardiac tubulin during chronic heart failure in the rat. Mol Cell Biochem. 2002;237:39–46

44. Sato H, Nagai T, Kuppuswamy D, Narishige T, Koide M, Menick DR, Cooper G, IV. Microtubule stabilization in pressure overload cardiac hypertrophy. Journal of Cell Biology. 1997;139:963–973

45. Nishimura S, Nagai S, Katoh M, Yamashita H, Saeki Y, Okada J, Hisada T, Nagai R, Sugiura S. Microtubules modulate the stiffness of cardiomyocytes against shear stress. Circ Res. 2006;98:81–87

46. Swiatlowska P, Sanchez-Alonso JL, Wright PT, Novak P, Gorelik J. Microtubules regulate cardiomyocyte transversal young’s modulus. Proc Natl Acad Sci U S A. 2020;117:2764–2766

47. Robison P, Caporizzo MA, Ahmadzadeh H, Bogush AI, Chen CY, Margulies KB, Shenoy VB, Prosser BL. Detyrosinated microtubules buckle and bear load in contracting cardiomyocytes. Science. 2016;352:aaf0659

48. Caporizzo MA, Chen CY, Bedi K, Margulies KB, Prosser BL. Microtubules increase diastolic stiffness in failing human cardiomyocytes and myocardium. Circulation. 2020;141:902–915

49. Grimes KM, Prasad V, McNamara JW. Supporting the heart: Functions of the cardiomyocyte’s non-sarcomeric cytoskeleton. J Mol Cell Cardiol. 2019;131:187–196

50. Wang B, Golemis EA, Kruh GD. Argbp2, a multiple src homology 3 domain-containing, arg/abl- interacting protein, is phosphorylated in v-abl-transformed cells and localized in stress fibers and cardiocyte z-disks. J Biol Chem. 1997;272:17542–17550

51. Luck K, Kim DK, Lambourne L, et al. A reference map of the human binary protein interactome.Nature. 2020;580:402–408

52. Soubeyran P, Barac A, Szymkiewicz I, Dikic I. Cbl-argbp2 complex mediates ubiquitination and degradation of c-abl. Biochem J. 2003;370:29–34

53. Cestra G, Toomre D, Chang S, De Camilli P. The abl/arg substrate argbp2/nargbp2 coordinates the function of multiple regulatory mechanisms converging on the actin cytoskeleton. Proc Natl Acad Sci U S A. 2005;102:1731–1736

54. Qiu Z, Cang Y, Goff SP. C-abl tyrosine kinase regulates cardiac growth and development. Proc Natl Acad Sci U S A. 2010;107:1136–1141

55. Zandy NL, Playford M, Pendergast AM. Abl tyrosine kinases regulate cell-cell adhesion through rho gtpases. Proc Natl Acad Sci U S A. 2007;104:17686–17691

56. Miller AL, Wang Y, Mooseker MS, Koleske AJ. The abl-related gene (arg) requires its f-actin- microtubule cross-linking activity to regulate lamellipodial dynamics during fibroblast adhesion. J Cell Biol. 2004;165:407–419

57. Colicelli J. Abl tyrosine kinases: Evolution of function, regulation, and specificity. Sci Signal. 2010;3:re6

58. Wang Y, Miller AL, Mooseker MS, Koleske AJ. The abl-related gene (arg) nonreceptor tyrosine kinase uses two f-actin-binding domains to bundle f-actin. Proc Natl Acad Sci U S A. 2001;98:14865–14870

59. Hu Y, Lyu W, Lowery LA, Koleske AJ. Regulation of mt dynamics via direct binding of an abl family kinase. J Cell Biol. 2019;218:3986–3997

60. Haglund K, Ivankovic-Dikic I, Shimokawa N, Kruh GD, Dikic I. Recruitment of pyk2 and cbl to lipid rafts mediates signals important for actin reorganization in growing neurites. J Cell Sci. 2004;117:2557–2568

61. Teckchandani AM, Birukova AA, Tar K, Verin AD, Tsygankov AY. The multidomain protooncogenic protein c-cbl binds to tubulin and stabilizes microtubules. Exp Cell Res. 2005;306:114–127

62. Hein MY, Hubner NC, Poser I, et al. A human interactome in three quantitative dimensions organized by stoichiometries and abundances. Cell. 2015;163:712–723

63. Sweet ES, Previtera ML, Fernández JR, Charych EI, Tseng C-Y, Kwon M, Starovoytov V, Zheng JQ, Firestein BL. Psd-95 alters microtubule dynamics via an association with eb3. The Journal of Neuroscience. 2011;31:1038

64. Glazier AA, Hafeez N, Mellacheruvu D, et al. Hsc70 is a chaperone for wild-type and mutant cardiac myosin binding protein c. JCI Insight. 2018;3

65. Sanger JM, Wang J, Gleason LM, Chowrashi P, Dube DK, Mittal B, Zhukareva V, Sanger JW. Arg/abl-binding protein, a z-body and z-band protein, binds sarcomeric, costameric, and signaling molecules. Cytoskeleton (Hoboken). 2010;67:808–823

66. Asimaki A, Tandri H, Huang H, et al. A new diagnostic test for arrhythmogenic right ventricular cardiomyopathy. N Engl J Med. 2009;360:1075–1084

67. Zhao L, Wang W, Huang S, et al. The rna binding protein sorbs2 suppresses metastatic colonization of ovarian cancer by stabilizing tumor-suppressive immunomodulatory transcripts. Genome Biol. 2018;19:35

68. Lv Q, Dong F, Zhou Y, Cai Z, Wang G. Rna-binding protein sorbs2 suppresses clear cell renal cell carcinoma metastasis by enhancing mtus1 mrna stability. Cell Death Dis. 2020;11:1056

69. Maurin ML, Labrune P, Brisset S, Le Lorc’h M, Pineau D, Castel C, Romana S, Tachdjian G. Molecular cytogenetic characterization of a 4p15.1-pter duplication and a 4q35.1-qter deletion in a recombinant of chromosome 4 pericentric inversion. Am J Med Genet A. 2009;149A:226–231

70. Molck MC, Simioni M, Paiva Vieira T, et al. Genomic imbalances in syndromic congenital heart disease. J Pediatr (Rio J). 2017;93:497–507

71. Liang F, Wang B, Geng J, et al. Sorbs2 is a genetic factor contributing to cardiac malformation of 4q deletion syndrome patients. Elife. 2021;10

72. Fu K, Nakano H, Morselli M, Chen T, Pappoe H, Nakano A, Pellegrini M. A temporal transcriptome and methylome in human embryonic stem cell-derived cardiomyocytes identifies novel regulators of early cardiac development. Epigenetics. 2018;13:1013–1026

73. Vigil-Garcia M, Demkes CJ, Eding JEC, et al. Gene expression profiling of hypertrophic cardiomyocytes identifies new players in pathological remodelling. Cardiovasc Res. 2021;117:1532–1545

74. Wales S, Hashemi S, Blais A, McDermott JC. Global mef2 target gene analysis in cardiac and skeletal muscle reveals novel regulation of dusp6 by p38mapk-mef2 signaling. Nucleic Acids Res. 2014;42:11349–11362

75. Dixon DM, Choi J, El-Ghazali A, Park SY, Roos KP, Jordan MC, Fishbein MC, Comai L, Reddy S. Loss of muscleblind-like 1 results in cardiac pathology and persistence of embryonic splice isoforms. Sci Rep. 2015;5:9042

76. Wang H, Bei Y, Shen S, et al. Mir-21-3p controls sepsis-associated cardiac dysfunction via regulating sorbs2. J Mol Cell Cardiol. 2016;94:43–53

77. Spengler RM, Zhang X, Cheng C, McLendon JM, Skeie JM, Johnson FL, Davidson BL, Boudreau RL. Elucidation of transcriptome-wide microrna binding sites in human cardiac tissues by ago2 hits- clip. Nucleic Acids Res. 2016;44:7120–7131

78. Ichikawa T, Kita M, Matsui TS, et al. Vinexin family (sorbs) proteins play different roles in stiffness-sensing and contractile force generation. J Cell Sci. 2017;130:3517–3531

